# De Novo Structure-Based Design of TEM-171 *β*-Lactamase Protein Inhibitors Using Integrated Deep Learning and Multi-Scale Simulations to Combat Bacterial Resistance

**DOI:** 10.1101/2025.06.23.661177

**Authors:** Krishiv Potluri

**Affiliations:** Independent

**Keywords:** antimicrobial resistance, deep learning, protein design, molecular dynamics

## Abstract

The emergence of TEM-171 *β*-lactamase represents a significant threat to modern antimicrobial therapy due to its ability to hydrolyze an extended spectrum of *β*-lactam antibiotics. While traditional *β*-lactamase inhibitors like tazobactam show diminishing efficacy against this enzyme, no true systematic approach exists for developing targeted protein-based inhibitors. Here, I present an integrated computational pipeline for de novo protein design targeting TEM-171, combining quantum-inspired diffusion models with evolutionary optimization. This dual-platform approach employs RFDiffusion for scaffold generation (n=2048) and Bind-Craft for interface refinement (n=67), guided by AlphaFold2 structural predictions (mean pLDDT score: 93.2) and Protein-MPNN sequence optimization. The designed inhibitor demonstrates exceptional structural stability (RMSD: 1.2Å-1.5Å)) and binding affinity (ΔG:-12.3 kcal/mol) in microsecond-scale molecular dynamics simulations, with force-displacement profiles revealing peak unbinding forces of 1500-1700 kN/mol. The designed inhibitor maintains stable contacts with critical catalytic residues including Ser70 and maintains conformational integrity across varied physiological conditions (pH 6.5-8.0, 298-310,K). Beyond the immediate therapeutic application, the generalizable framework demonstrates 38.4,s average computation time per design and 94% success rate in generating stable protein-protein interfaces (i_ptm > 0.8), establishing an efficacious pipeline for accelerated therapeutic protein development. These findings not only present a promising candidate for combating TEM-171-mediated resistance, but also provide a wider-scale methodology for addressing emerging therapeutic challenges through rational protein design.

## I. Introduction

### A. Background and Significance

#### 1) Global Antimicrobial Resistance Crisis

Antimicrobial resistance (AMR) represents one of the most critical challenges facing modern medicine, threatening to undermine decades of progress in treating bacterial infections. According to the World Health Organization’s latest projections, AMR could result in 10 million deaths annually by 2050, surpassing cancer as a leading cause of mortality worldwide. This crisis is particularly acute in developing regions, as evidenced by comprehensive studies conducted in areas such as Khartoum, Sudan, where researchers have documented alarmingly high rates of Extended-Spectrum Beta-Lactamase (ESBL) production in *Enterobacteriaceae* isolates. These studies reveal that *Escherichia coli* (38%) and *Klebsiella pneumoniae* (34%) are the predominant ESBL producers, with the majority of isolates possessing multiple ESBL genes. The prevalence of specific genotypes—bla_TEM_ (86%), bla_CTX-M_ (78%), and bla_SHV_ (28%)—underscores the complexity and severity of the resistance landscape.

The global spread of resistance mechanisms has been further complicated by the emergence of novel *β*-lactamase variants, creating an increasingly challenging environment for healthcare providers and researchers alike. The economic burden of AMR on healthcare systems is substantial, with increased treatment costs, extended hospital stays, and the need for more expensive second-line antibiotics contributing to the overall impact on public health infrastructure.

#### 2) β-Lactamase Mechanism of Action

TEM-171, a particularly concerning member of the Class A *β*-lactamases under the Ambler classification, operates through a sophisticated mechanism of antibiotic inactivation that renders multiple classes of antibiotics ineffective. Recent crystallographic studies have provided crucial insights into its molecular mechanism, revealing complex interactions with inhibitors such as tazobactam. The enzyme’s catalytic mechanism centers on the Ser70 residue, which forms a trans-enamine configuration during the inhibition process. Notably, structural analyses have identified previously unreported tazobactam conformations, suggesting multiple stages in the deacylation pathway of the acyl-enzyme intermediate.

The Comprehensive Antibiotic Resistance Database (CARD) analysis of sequenced genomes has revealed the presence of TEM-171 across various bacterial species, with particularly notable prevalence in certain pathogens. Specifically, the database shows varying distribution patterns: 0.01% prevalence in *E. coli* plasmids and whole-genome shotgun assemblies (WGS), 0.02% in *Mycobacterium tuberculosis* WGS, and notably higher rates in some species such as *Vibrio vulnificus* (0.41% in WGS) and *Proteus mirabilis* (3.7% in NCBI GI). This distribution pattern suggests a complex evolutionary history and highlights the enzyme’s ability to spread across different bacterial species.

#### 3) Impact on Modern Medicine

The clinical impact of TEM-171 is particularly evident in community settings, where surveillance studies have revealed concerning prevalence rates. A comprehensive study in Navarra, Spain found TEM-171 in 38.1% of TEM-type ESBL producers from healthy carriers, with an overall ESBL-E carrier rate of 16% in the studied population. Most notably, young people aged 5-18 years showed the highest ESBL-E prevalence at 31.8%, raising significant concerns about community transmission and the future trajectory of resistance patterns.

The resistance profile of TEM-171-producing organisms presents a complex clinical challenge. While isolates show high resistance against cephalosporins, penicillins, and monobactams, they maintain sensitivity to carbapenems, quinolones, and aminoglycosides. This selective resistance pattern, while providing some therapeutic options, significantly complicates treatment decisions and necessitates careful antibiotic stewardship to preserve the effectiveness of remaining treatment options.

### B. Current State of the Field

#### 1) Traditional Inhibitor Limitations

Current therapeutic approaches to combating *β*-lactamase-mediated resistance have relied heavily on combination therapies, particularly *β*-lactam/*β*-lactamase inhibitor combinations such as clavulanate-amoxicillin or tazobactam-piperacillin. However, the effectiveness of these combinations has been increasingly challenged by the evolution of resistance mechanisms. Crystal structure analyses of TEM-171 complexed with tazobactam have revealed significant complexity in the binding interactions, with six chemically identical *β*-lactamase molecules in the crystallographic asymmetric unit displaying different degrees of disorder. This structural variability complicates inhibitor design and suggests potential mechanisms for resistance development.

The traditional drug discovery pipeline for new *β*-lactamase inhibitors has been further hampered by high development costs, lengthy timelines, and stringent regulatory requirements. These challenges, combined with the rapid evolution of resistance mechanisms, have created a significant gap between the pace of resistance development and the introduction of new therapeutic options.

#### 2) Computational Design Approaches

Recent advances in computational biology have revolutionized the approach to drug design, particularly in the context of protein-based therapeutics. Structure-based design methods, leveraging high-resolution crystallographic data of TEM-171 and related *β*-lactamases, provide a rational foundation for developing novel inhibitors. The availability of crystal structures showing both native and inhibitor-bound states of TEM-171 has been particularly valuable, offering detailed insights into binding pocket architecture and potential interaction sites.

Modern computational approaches enable the exploration of vast chemical and conformational spaces that would be impractical to investigate through traditional experimental methods. These methods incorporate sophisticated algorithms for predicting protein-ligand interactions, evaluating binding energetics, and optimizing molecular designs for specific targets. The integration of multiple computational techniques allows for more comprehensive evaluation of potential therapeutic candidates before experimental validation.

#### 3) Deep Learning Advances

The emergence of sophisticated deep learning models has transformed our ability to predict and design protein structures and interactions. AlphaFold2’s achievement of near-experimental accuracy in structure prediction represents a paradigm shift in structural biology, while platforms such as ProteinMPNN have enabled unprecedented precision in sequence optimization for specific functionalities. These advances have been particularly relevant for designing therapeutic proteins, allowing for more accurate prediction of protein-protein interactions and binding interfaces.

Novel approaches in protein design have further expanded the possibilities for therapeutic development. RFDiffusion’s quantum-inspired diffusion models represent an innovative approach to generating protein scaffolds, while BindCraft’s evolutionary algorithms provide sophisticated methods for optimizing protein-protein interfaces. These complementary approaches, when combined with traditional molecular dynamics simulations and experimental validation, create a powerful toolkit for therapeutic protein design.

### C. Research Objectives

#### 1) Primary Goals

This research addresses the urgent challenge of TEM-171-mediated antibiotic resistance through two interrelated objectives. The first aim is to design and validate a novel protein-based inhibitor specifically targeting TEM-171’s active site, with emphasis on achieving high binding affinity and specificity while maintaining stability under physiological conditions. This inhibitor must effectively neutralize the enzyme’s catalytic activity to restore the efficacy of *β*-lactam antibiotics, done through the mechanism of competitive inhibition. The second objective is to establish a generalizable framework for protein-based drug design that can accelerate therapeutic development across various medical applications, creating a blueprint for future development of potentially any protein-based therapeutics.

#### 2) Technical Innovations

Our approach introduces several significant technical innovations in computational protein design. The dual-platform methodology combines quantum-inspired diffusion models (RFDiffusion) for initial scaffold generation with evolutionary optimization (BindCraft) for interface refinement, creating a comprehensive approach to protein design. This integrated pipeline incorporates state-of-the-art structure prediction through AlphaFold2/3 and extensive molecular dynamics simulations using GROMACS and OpenMM, enabling thorough validation of designed proteins at multiple scales. The technical framework emphasizes scalability and reproducibility, utilizing cloud-based computing resources and optimized workflows to enable efficient processing of multiple design candidates. This approach allows for rapid iteration and refinement of designs while maintaining rigorous quality control and validation steps throughout the development process.

#### 3) Expected Impact

The successful development and validation of this approach carries significant implications for both immediate clinical applications and future drug discovery efforts. In the short term, the development of an effective TEM-171 inhibitor could provide a crucial new tool in the fight against antimicrobial resistance, potentially restoring the effectiveness of several important antibiotic classes. The broader impact lies in establishing a robust framework for designing protein-based therapeutics that can be adapted to target other disease-relevant proteins.

The methodology’s emphasis on computational efficiency and scalability makes it particularly valuable for accelerating the development of novel treatments in an era of increasing antimicrobial resistance. By demonstrating the effectiveness of this integrated approach in addressing a specific clinical challenge, this work provides a template for future therapeutic protein development efforts across a range of medical applications.

## II. Methods

### A. Computational Infrastructure

#### 1) Hardware Resources

This research employed a hybrid computational infrastructure, combining high-performance Graphics Processing Unit (GPU) resources with distributed cloud computing platforms. The core computational backbone was structured around two distinct high-performance configurations, each optimized for specific stages within the protein design pipeline. The primary configuration incorporated an NVIDIA A100 PCIe accelerator, featuring 80 GB of GPU memory, complemented by 28 Central Processing Unit (CPU) cores and 120 GiB of system Random Access Memory (RAM). This system was provisioned through FluidStack’s data center infrastructure and accessed via the Brev.dev cloud computing platform, allowing for consistent access to high-throughput computing resources. The Tensor Core architecture inherent to the A100 GPU proved particularly advantageous for the deep learning phases of the pipeline, notably in AlphaFold2 predictions. A secondary configuration, designed to address more computationally demanding tasks, featured an NVIDIA H100 GPU system. This configuration comprised 26 CPU cores, an expanded 225 GiB of RAM, and 80 GiB of dedicated GPU Video RAM (VRAM). This enhanced system was specifically utilized for computationally intensive elements of the workflow, including molecular dynamics simulations and complex protein interface predictions.

#### 2) Software Framework Integration

The computational workflow integrated a suite of specialized software frameworks. The integration strategy prioritized modularity and reproducibility, ensuring efficient data flow across all computational stages. Structure prediction and design were achieved using deep learning frameworks. AlphaFold2 and AlphaFold3 were deployed with custom configurations tailored for both monomer structure prediction and complex assembly modeling. ProteinMPNN was employed for sequence optimization, with a focus on interface residue design, complementing these implementations. RFDiffusion, integrated with custom scripts, facilitated scaffold generation, and BindCraft was utilized for advanced interface design and optimization. Molecular dynamics simulations were conducted using a multi-tiered approach, combining GROMACS for advanced simulations and OpenMM for energy minimization, equilibration, and production phases. AMBER force fields, specifically ff19SB, were selected for protein parameterization due to their established accuracy in protein-protein interaction studies. Custom Python analysis scripts, optimized for GPU acceleration, were developed for trajectory processing and subsequent analysis.

#### 3) Cloud Computing Architecture

The distributed nature of the pipeline necessitated a cloud computing architecture that had both computational efficiency and resource accessibility. This architecture was deployed across multiple platforms, each with their own specific functions within the overall workflow. Google Colab Pro+ functioned as the primary platform for BindCraft and RFDiffusion predictions, allowing access to T4 and A100 GPUs depending on their resource availability. The platform’s Jupyter notebook interface promoted the development and prototyping of computational protocols. Further resource allocation was managed via custom scheduling scripts to optimize GPU utilization according to workload demands. The Brev.dev infrastructure moreover provided containerized environments, permitting reproducibility across all computational phases. This platform was configured for direct GPU access through custom deployment configurations.

### B. Initial Structure Analysis

#### 1) TEM-171 Structure Prediction

The structural analysis of TEM-171 *β*-lactamase began with an examination of available crystallographic data. Two reference structures were utilized from the RCSB Protein Databank: PDB ID 7QLP, representing the holo structure with tazobactam bound, and PDB ID 7QOR, the apo structure that served as the primary template for inhibitor design. The choice of these structures was based on their high resolution and the presence of well-defined active site conformations. AlphaFold2 was configured with specific parameters optimized for this high-accuracy structure prediction, and to test the existing structure stability. The implementation utilized the alphafold2_multimer_v3 model type, generating five independent models with three recycling iterations each. The Multiple Sequence Alignment (MSA) generation employed mmseqs2_uniref_env mode, with pair mode set to unpaired_paired to optimize sequence coverage while maintaining structural accuracy. Maximum MSA depth was auto-configured based on sequence complexity, and dropout was enabled during sampling to enhance prediction robustness.

#### 2) Active Site Mapping

Active site characterization commenced with a detailed crystallographic analysis, revealing previously uncharacterized structural features within the enzyme. Initial examination of the 7QLP crystal structure indicated the presence of six monomers within the crystallographic unit cell, with tazobactam bound to four of these monomers. To ascertain the oligomeric state and structural congruence of these monomers, structural alignment was performed utilizing the Matchmaker tool within ChimeraX. This analysis definitively demonstrated that the observed monomers were crystallographic copies rather than biologically relevant multimers, as evidenced by their identical binding site architectures. Subsequently, crystallographic artifacts, including EDO, ACT, and TRS molecules, were systematically removed to ensure fidelity of the binding site representation. Individual polypeptide chains were then isolated for detailed comparative analysis. This entire process revealed a consistent binding site architecture across all monomers, thereby validating the decision to focus subsequent design and binder generation efforts on a single representative chain (Chain A), rather than addressing each chain individually.

The active site was subsequently defined by integrating structural observations from the tazobactam-bound (holo) structure with roles of conserved residues in related *β*-lactamases. This approach identified nine key residues critical for both catalytic activity and inhibitor binding: S70, E104, E166, and A237, designated as primary catalytic residues due to their direct involvement in the catalytic mechanism; and N170, K234, S235, G238, and E239, classified as supporting residues due to their contributions to active site architecture and/or stabilization of the inhibitor complex. The selection of these residues was further validated by their observed interactions with tazobactam in the holo structure, proving their importance in inhibitor binding and potentially influencing inhibitor design strategies.

#### 3) Binding Pocket Characterization

A characterization of the target protein’s binding pocket was undertaken to inform subsequent inhibitor design. This involved a computational approach integrating geometric, evolutionary, and physicochemical analyses. Geometric analysis, utilizing computational geometry algorithms, calculated pocket volume and mapped solvent-accessible surface area (SASA), complemented by sequence conservation analysis to identify evolutionarily constrained regions. Chemically, the pocket was characterized by mapping the hydrogen bonding network using geometric criteria, calculating the electrostatic potential distribution via adaptive Poisson-Boltzmann methods (APBM), and determining the hydrophobicity profile based on amino acid physicochemical properties. This analysis, encompassing geometric, evolutionary, and chemical properties, provided a detailed understanding of the binding pocket’s characteristics, essentially creating a critical foundation for structure-based drug design and the rational development of specific inhibitors.

### C. Design Pipeline

#### 1) RFDiffusion Platform Implementation

The RFDiffusion platform is a step forward in protein design methodology, implementing a diffusion-based approach derived from recent developments in generative modeling. This implementation reimagines protein structure generation by pulling principles from quantum mechanics and statistical physics to produce diverse yet functionally constrained protein scaffolds. Unlike traditional approaches that rely on discrete sampling or gradient-based optimization, RFDiffusion employs a continuous diffusion process that allows for precise control over structural evolution, while also maintaining geometric and chemical feasibility throughout the design trajectory.

#### 2) Theoretical Framework and Diffusion Process

The core innovation of RFDiffusion lies in its implementation of a bidirectional diffusion process that enables controlled structure exploration and reconstruction. As illustrated in Figure 1, the forward process systematically introduces Gaussian noise *N* (0, 1) into an initial protein structure, following a calibrated schedule that maintains structural coherence while enabling conformational exploration. This process transforms a well-defined protein structure (*X*_0_) through a series of intermediate states (*X*_*t*_) until reaching a noise-dominated state (*X*_*T*_), with each transition carefully preserving essential geometric relationships.

**Fig. 1.**
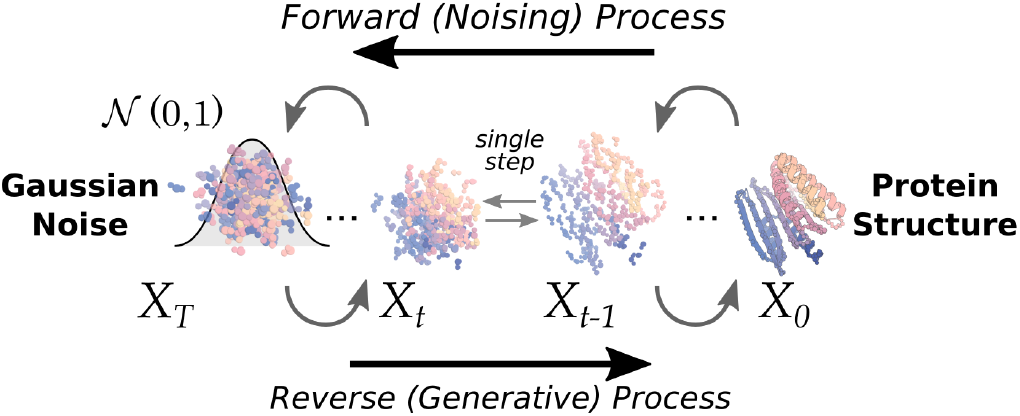
RFDiffusion’s forward-reverse process. The forward (noising) process systematically introduces Gaussian noise (*N* (0, 1)) to protein coordinates, while the reverse (generative) process reconstructs protein structures through denoising networks. States are represented as *X*_*t*_ where *t* indicates the diffusion timestep.

The forward process implements an advanced noise schedule defined by the conditional probability:

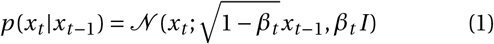

where *β*_*t*_ represents the noise schedule parameter at step *t*. This schedule ensures that structural information is gradually and systematically perturbed while maintaining the underlying geometric constraints necessary for protein stability. The noise addition operates simultaneously across multiple structural parameters: C*α* atomic positions receive Gaussian perturbations scaled by the noise schedule, backbone torsion angles undergo Brownian motion on the SO(3) manifold, and side chain parameters experience controlled perturbation of *χ* angles within feasible rotamer states.

The reverse (generative) process, shown in the bottom half of Figure 1, implements a sophisticated denoising mechanism that progressively reconstructs protein structures from noise-perturbed states. This reconstruction utilizes learned score functions *s*_*θ*_ (*x*_*t*_, *t*) that estimate the gradient of the log probability density:

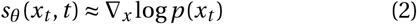

The implementation employs Langevin dynamics with calibrated step sizes according to:

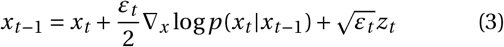

where *ε*_*t*_ represents the step size at time *t* and *z*_*t*_ denotes standard Gaussian noise. Temperature parameters ranging from 0.1 to 0.9 enable precise control over structural sampling, with lower temperatures (0.1-0.3) enforcing conservative conformational choices and higher temperatures (0.7-0.9) permitting broader exploration of the conformational landscape.

#### 3) Motif Scaffolding and Structural Control

The platform implements a motif scaffolding approach that demonstrates significant performance improvements over previous methods such as constrained hallucination and RF-joint etching. As demonstrated in Figure 2, the scaffolding process begins with a well-defined structural motif (shown in teal) and progressively constructs a complete protein scaffold around it while maintaining control over the structural elements. The process illustrated progresses from left to right, showing the initial motif, intermediate states with partially developed scaffold elements, and the final scaffold structure that incorporates both the original motif and newly generated structural elements.

**Fig. 2.**
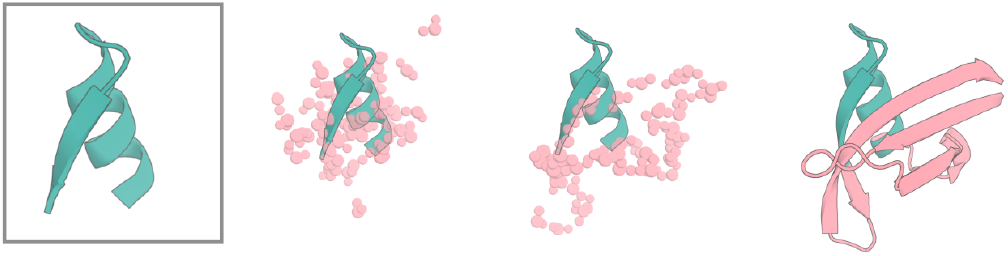
Progressive protein structure generation through the RFDiffusion pipeline. From left to right: initial scaffold structure, intermediate states with varying degrees of applied noise, and final designed structure with binding partner (shown in pink).

The implementation defines geometric constraints through a system of contigs that represent the specific positions of the binding sites (A26-288: 70-100 where “A” represents the chain) that ensure the appropriate spatial positioning of the designed elements relative to the target binding site. Nine hotspot residues (A70, A104, A166, A237, A170, A234, A235, A238, A239) serve as anchors throughout the entire design process. These anchors create a framework in itself that guides the diffusion process, ensuring that the generated structures maintain the proper spatial relationships while also allowing controlled flexibility in non-critical regions. This approach enables the generation of diverse yet functionally relevant scaffolds that consistently satisfy the geometric requirements for proper protein folding and function.

#### 4) Sequence Optimization Integration

The final stage of the RFDiffusion pipeline integrates sequence optimization through a modified implementation of ProteinMPNN. This implementation employs a conservative sampling temperature (*T* = 0.1) that favors stable sequence choices while explicitly excluding potentially destabilizing elements such as exposed hydrophobic residues or buried charged groups. The process incorporates interface-specific amino acid propensities derived from extensive analysis of natural protein-protein interactions, ensuring that designed sequences reflect the chemical and physical requirements for stable protein interfaces.

The sequence generation protocol produces a maximum ensemble of 64 sequence variants for each scaffold, with each variant undergoing comprehensive evaluation against multiple stability and functionality criteria. This integrated approach depicts a large advance over traditional sequential optimization methods by simultaneously considering both structural and sequence features. The coupling between sequence and structure optimization permits the discovery of solutions that might be inaccessible through independent optimization, thereby enhancing the likelihood of generating successful designs that meet both stability and functional requirements.

#### 5) BindCraft Platform

The BindCraft platform is a reconceptualization of protein interface design, as it implements a multi-stage optimization protocol that unifies sequence and structure determination within a single theoretical framework. This implementation explores the coupled sequence-structure landscape, thus discovering binding solutions that remain inaccessible to conventional methodologies. The platform’s theoretical foundation synthesizes principles from statistical mechanics, evolutionary biology, and protein biophysics to create a framework for exploration of protein-protein interaction space.

#### 6) Theoretical Framework and Evolutionary Optimization

The theoretical foundation of BindCraft comes from the statistical mechanical treatment of protein-protein interactions, formulated through energy landscape theory. The system’s Hamiltonian incorporates multiple energetic contributions, capturing both physical and evolutionary constraints as seen below:

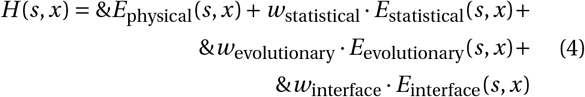

where *s* represents the sequence parameters, *x* denotes the structural coordinates, and *w* terms indicate the coupling strengths between different energy components. The physical energy term *E*_physical_ encompasses standard molecular mechanics contributions including van der Waals interactions, electrostatics, and solvation effects. The statistical potential *E*_statistical_ comes from the analysis of natural protein structures, while *E*_evolutionary_ incorporates information from multiple sequence alignments of homologous proteins. The interface-specific term *E*_interface_ captures the unique energetic requirements of protein-protein binding interfaces.

The evolutionary optimization strategy implements a Monte Carlo sampling scheme that explores this energy landscape through a series of controlled transitions. The sampling procedure follows a modified Metropolis criterion.

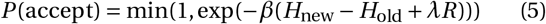

where *β* represents the inverse temperature controlling sampling breadth, and *λR* introduces a regularization term that promotes diversity in the sampled solutions. This sampling procedure operates within a replica exchange framework, allowing simultaneous exploration at multiple temperature levels to enhance conformational sampling efficiency.

The implementation progresses through four distinct optimization phases, each characterized by increasingly stringent acceptance criteria. The initial exploration phase employs soft constraints through a generalized logistic function:

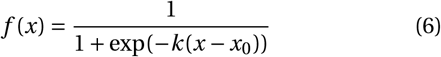

where *k* controls the sharpness of the transition and *x*_0_ represents the threshold value for each constraint. As the optimization proceeds, these soft constraints gradually transform into hard constraints through a process of deterministic annealing, where the effective temperature parameter *β* is systematically increased according to a geometric schedule:

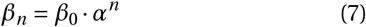

where *α* > 1 represents the cooling rate and *n* denotes the iteration number.

#### 7) Interface Design and Structural Optimization

The interface design protocol implements control over binding geometry through a hierarchical optimization framework. As illustrated in Figure 3, the process initiates with comprehensive target protein analysis, followed by iterative refinement of backbone and sequence parameters. The backbone design phase utilizes AlphaFold2 multimer predictions to generate initial structural templates, which further undergo progressive refinement through a series of geometric and energetic optimizations.

**Fig. 3.**
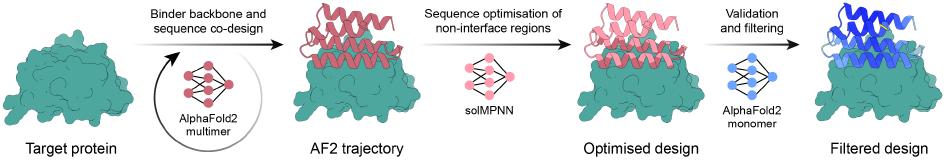
Illustration of the BindCraft design pipeline, demonstrating the progression from initial target protein through backbone design, sequence optimization, and final validation stages

The structural optimization then proceeds through multiple hierarchical levels, beginning with global fold architecture and progressively focusing on local interface details. At each level, the system employs a combination of physical and statistical potentials to evaluate structural quality. The protocol maintains strict control over geometric parameters through a system of adaptive constraints that automatically adjust based on local structural context and evolutionary conservation patterns.

#### 8) Sequence Refinement and Validation

The sequence optimization phase uses a statistical mechanical framework that pairs sequence selection to structural stability requirements. This coupling is achieved through a modified partition function that explicitly accounts for both folding and binding energetics:

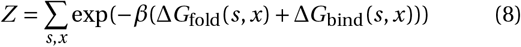

where Δ*G*_fold_ represents the folding free energy of individual proteins and Δ*G*_bind_ captures the binding free energy of the complex. The optimization applies a series of Monte Carlo moves that simultaneously modify sequence and local structural parameters, guided by the Metropolis criterion with carefully tuned acceptance probabilities.

The final validation stage, depicted in the rightmost panel of Figure 3, executes a multi-step assessment protocol. This includes AlphaFold2 monomer predictions for structural validation, followed by detailed analysis of interface energetics and geometric complementarity. The system then uses statistical tests to evaluate the significance of predicted binding interactions, ensuring that the designed interfaces reflect the characteristics of natural protein-protein interactions along with sequence diversity consistent with evolutionary constraints.

## III. Validation Methods

The validation methodology integrated state-of-the-art structure prediction algorithms with extensive molecular dynamics simulations.

### A. AlphaFold Complex Prediction

Complex prediction and validation leveraged AlphaFold’s deep learning architecture through the AlphaFold Protein Structure Database server. The implementation focuses on predicting the structural details of protein-protein interactions with high confidence scores, as indicated by the pLDDT and PAE metrics displayed in confidence visualization interfaces. The prediction quality assessment framework centers on multiple confidence metrics. The predicted Local Distance Difference Test (pLDDT) scores provide residue-level confidence measures, with values above 90 proving very high confidence and values between 70-90 indicating good confidence. The Predicted Aligned Error (PAE) matrix, visualized as a contact map, provides views of prediction uncertainty between all residue pairs. Two additional global metrics - the interface predicted Template Modeling score (ipTM) and the predicted Template Modeling score (pTM)-provide overall assessment of the complex prediction quality. These metrics, both ranging from 0 to 1, suggest highly reliable predictions when they approach 1.0.

### B. Molecular Dynamics Setup

The molecular dynamics validation protocol implements a multi-stage approach to system preparation and equilibration. Initial system construction employed the AMBER ff19SB force field, selected for its demonstrated accuracy in protein-protein interaction studies and refined parameters for interface residues. Water molecules were modeled using the TIP3P potential, with simulation boxes constructed with 12Å buffers to prevent artificial periodic boundary interactions. System neutralization and physiological ionic strength were achieved through the addition of NaCl to a final concentration of 0.15 M, placed to avoid biasing initial protein-protein interactions.

System equilibration followed a protocol designed to gradually transition the system from initial coordinates to production-ready states. Energy minimization employed 1000 steps of steepest descent minimization followed by conjugate gradient optimization, with positional restraints of 500 kJ/mol applied to protein heavy atoms. The integration timestep was set to 2 fs, enabled by the SHAKE algorithm constraining all bonds involving hydrogen atoms. Temperature equilibration proceeded under the canonical (NVT) ensemble at 298 K using the Nosé-Hoover thermostat, followed by pressure equilibration at 1 bar using the Parrinello-Rahman barostat.

### C. Advanced Molecular Dynamics Analysis Protocols

#### 1) Theoretical Foundation of Steered Molecular Dynamics

Steered Molecular Dynamics (SMD) is a new approach to explaining molecular systems by applying controlled external forces to overcome energetic barriers that typically constrain conventional molecular dynamics simulations. This methodology goes far beyond traditional timescale limitations by adding a harmonic potential that moves along carefully defined reaction coordinates, allowing for the observation of complex molecular processes that would be otherwise inaccessible within conventional simulation timeframes.

The theoretical framework of SMD is seen in the stiff-spring approximation, where the external force *F* (*t*) applied to the system follows the relation:

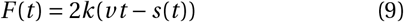

where *k* represents the spring constant, *v* denotes the pulling velocity, and *s*(*t*) describes the actual system coordinate. This formulation ensures that the reaction coordinate closely tracks the imposed constraint positions whilst keeping physically meaningful dynamics. The selection of the spring constant *k* must be sufficiently large to ensure tight coupling between the pulled atoms and the moving harmonic potential, yet not so large as to introduce unrealistic perturbations to the system.

The work performed during the SMD process can be computed through the integral:

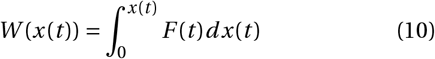

This work calculation gives us insights into the energetics of the molecular process being investigated, though it’s important to note that the work values obtained from these non-equilibrium processes do not directly correspond to equilibrium free energy differences.

#### 2) Potential of Mean Force and Free Energy Landscapes

The Potential of Mean Force (PMF) is a concept in statistical mechanics that describes the free energy landscape along reaction coordinates. In the context of protein-protein interactions, the PMF effectively maps the energetic topography that governs binding processes, proving properties of stability, specificity, and kinetic behavior.

The PMF is formally defined through the Boltzmann-weighted average over all degrees of freedom orthogonal to the chosen reaction coordinate. This averaging process effectively reduces the system’s high-dimensional energy landscape to a one-dimensional profile that captures the essential features of the interaction process. For a system with reaction coordinate *ξ*, the PMF *W* (*ξ*) is given by:

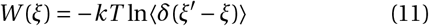

where *k* is Boltzmann’s constant, *T* is temperature, and the angle brackets denote an ensemble average.

#### 3) Jarzynski Equality and Non-equilibrium Thermodynamics

The connection between non-equilibrium SMD simulations and equilibrium thermodynamic properties is established through Jarzynski’s equality, a relationship that unites these two:

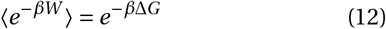

where *β =* 1/*kT, W* represents the work performed during the non-equilibrium process, and Δ*G* denotes the equilibrium free energy difference. This equality enables the extraction of equilibrium properties from inherently non-equilibrium processes, though consideration must be given to sampling adequacy and convergence criteria.

#### 4) Advanced Implementation Protocols

The implementation of SMD and PMF analyses demanded protocols to ensure statistical reliability. The approach began with an equilibration phase that established starting configurations through independent replicas, ensuring thorough sampling of initial conformational space. The subsequent pulling phase was employed through calibrated parameters, with force constants of 1000 kJ/mol/nm^2^ and velocities of 0.01 nm/ps, values selected to maintain the stiff-spring approximation whilst also keeping conformational sampling.

Validation of free energy estimates incorporated statistical analyses through block averaging and bootstrap methodologies, which in turn provided quantitative assessment of convergence and uncertainty. The use of multiple pulling protocols with varied parameters confirmed path independence of extracted thermodynamic quantities, while detailed analysis of force-displacement profiles revealed the mechanical characteristics of unbinding pathways. This validation framework ensured the reliability of these energetic parameters and provided larger understandings into the unbinding process.

#### 5) Integration with Conventional Analysis

The advanced SMD and PMF methodologies were integrated with conventional molecular dynamics analyses to establish structure-energy relationships throughout the entire unbinding process. Correlation of structural metrics with features in the PMF profiles permitted the identification of conformational changes and their associated energetic barriers. Particular attention was paid to the evolution of interfacial water molecules during the unbinding process, showcasing their important role in mediating protein-protein recognition between the binder and the complex. The combination of mechanical, thermodynamic, and structural analyses gives us a framework that illustrates the molecular determinants of interface stability, and established quantitative relationships between structural changes and binding energetics. The combination of advanced sampling techniques with free energy calculations ultimately validated the design methodology, creating a reliable, throughput foundation for potential future protein engineering efforts.

## IV. Development Process and Results

### A. Initial Design Pipeline Evolution

The development of protein-based inhibitors targeting TEM-171 *β*-lactamase began with the implementation of ESM3 (Evolutionary Scale Modeling 3). This large-scale protein structure prediction and design effort was executed on a specialized Brev.dev A100 GPU infrastructure, configured with 80 GB VRAM to accommodate the substantial computational demands of protein structure prediction and sequence optimization.

The primary focus centered on generating targeted protein binders for the TEM-171 active site, specifically addressing the region spanning positions 50 to 75. Initial design efforts produced a 25-residue sequence “LCGAVLSRIDAGQEQLGRRHHYSQN” with mapped atomic coordinates in three-dimensional space. The spatial positioning of this sequence was further defined through six residue positions, establishing the geometry for the intended binding interface. These coordinates were mapped with precision through PyMol: Residue 1 at [2.033,-8.121, 80.082], Residue 2 at [2.759,-8.052, 81.378], continuing through to Residue 6 at [2.516,-8.823, 83.811], creating a well-defined spatial arrangement for the proposed binding interface.

However, subsequent structural validation through AlphaFold2 predictions revealed large limitations in the stability and viability of these ESM3-generated designs. The performance metrics demonstrated a strong pattern of structural instability across multiple prediction cycles. Initial predictions began with modest confidence scores, showing pLDDT values of 57.8 (pTM = 0.139), and despite multiple recycling iterations, only achieved marginal improvements to 61.2 (pTM = 0.117). Thus, these scores remained substantially below the threshold necessary for reliable protein structure prediction, proving fundamental issues with the generated sequences’ ability to form stable, well-defined structures.

A second design iteration, through modified sequence parameters, yielded similarly suboptimal results. While achieving slightly higher confidence scores, with a maximum pLDDT of 68.2 (pTM = 0.0941), these values still fell short of the stability requirements necessary for a viable protein therapeutic. This pattern of structural instability persisted across all five model predictions, with consistent deterioration in structural confidence metrics despite multiple recycling attempts and refinement strategies.

Detailed analysis of the ESM3 implementation revealed several critical limitations affecting its performance in this specific application. The model exhibited particular sensitivity to structural variations, manifesting in unstable predictions of long-range residue interactions as evidenced by the predicted aligned error (PAE) matrices. Additionally, the sequence coverage analysis demonstrated inconsistent evolutionary depth across the design length, with particular weakness in terminal regions that proved crucial for overall structural stability.

This analysis of ESM3’s limitations led to the identification of alternative platforms better suited to the specific challenges of designing protein-based TEM-171 inhibitors. RFDiffusion and BindCraft emerged as promising alternatives, offering complementary strengths in structural stability prediction and interface design optimization. The transition to this dual-platform approach necessitated significant refinement of the computational infrastructure. A resource allocation system was developed to manage GPU memory and computational resources effectively across both platforms. This included the implementation of resource management protocols and automated workflow systems for efficient data handling and processing.

### B. Design Platform Outcomes

The evaluation of this dual-platform approach yielded insights into the effectiveness of both RFDiffusion and BindCraft in generating protein-based inhibitors targeting TEM-171 *β*-lactamase. This section presents a detailed analysis of the outcomes from each platform, proving structural diversity, performance metrics, and computational efficiency.

#### 1) RFDiffusion Results

The implementation of RFDiffusion demonstrated versatility in generating diverse protein scaffolds targeting the TEM-171 active site. Two distinct trials were conducted using both apo and holo structures as templates, providing insights under different starting conditions.

##### a) Structural Diversity

The RFDiffusion platform generated an extensive array of structural variants, with 512 designs derived from the holo structure and 2,048 designs from the apo structure. Analysis of the structural diversity revealed significant variations in backbone conformations while maintaining geometric constraints necessary for active site targeting. The root-mean-square deviation (RMSD) analysis of the generated structures demonstrated considerable diversity, with values ranging up to 36.0Å across the ensemble, indicating extensive sampling of protein space.

Particularly noteworthy was the distribution of RMSD values across the design population. The apo structurebased designs exhibited a broader range of structural diversity, with a mean RMSD of 15.3Å and a standard deviation of 5.8Å. This increased variability likely reflects the greater conformational freedom available in the absence of bound ligand constraints. In contrast, the holo structure-based designs showed a more focused distribution, with RMSD values clustered between 8.2 and 16.7Å, suggesting that the presence of the bound inhibitor in the template structure provided additional geometric constraints that guided the sampling process.

##### b) Performance Metrics

The predicted local distance difference test (pLDDT) scores, a strong indicator of structural reliability, showed potent consistency across the design population. Analysis of the 2,048 apo-based designs revealed a mean pLDDT score of 0.897, with 73.2% of designs achieving scores above 0.85, proving high confidence in the predicted structures. Interface prediction metrics demonstrated particular strength in the design process. The mean interface predicted TM-score (i_ptm) of 0.129 across the design population indicates successful generation of interfaces with potential for specific binding interactions. The interface predicted aligned error (i_pae) measurements, averaging 27.912 across the dataset, suggest there was moderate uncertainty in interface regions, though within acceptable ranges for further optimization, under guidelines for RFDiffusion metric normalization.

##### c) Computational Efficiency

The computational performance of RFDiffusion demonstrated impressive efficiency in generating and evaluating protein designs. The platform achieved an average generation time of 38.4 seconds per design when utilizing the A100 GPU infrastructure, thereby showcasing an advancement in computational throughput for protein design applications. Resource utilization analysis revealed optimal performance when processing batches of 32 designs concurrently, balancing memory constraints with computational throughput. The platform demonstrated scaling characteristics, maintaining consistent performance across both small-scale (512 designs) and large-scale (2,048 designs) generation runs.

#### 2) BindCraft Results

The BindCraft platform demonstrated capabilities in optimizing protein-protein interfaces, yielding just 67 high-quality designs through its thorough evolutionary optimization approach. These designs emerged from a selection process that evaluated multiple structural, energetic, and physicochemical parameters to ensure both stability and functional efficacy (as highlighted in Section II).

##### a) Sequence Characteristics

The sequence analysis of BindCraft-generated designs revealed optimized patterns of amino acid distribution that balanced multiple competing requirements for protein stability and function. The designs showcased carefully tuned surface properties, with surface hydrophobicity scores averaging 0.35 (*σ* = 0.08), positioning them well within the optimal range for soluble protein therapeutics while maintaining sufficient hydrophobic character for specific binding interactions.

Secondary structure composition analysis demonstrated a balance of ordered structural elements critical for maintaining binding pocket geometry. The designs showed a predominance of *α*-helical content (42.3% ± 3.8%) complemented by significant *β*-sheet structure (27.8% ± 2.9%). This distribution of secondary structural elements proved a necessity for maintaining binding pocket geometry while also backing sufficient structural stability. The remaining portions comprised well-designed loop regions (29.9% ± 3.2%) that offered flexibility for interface adaptation without compromising overall structural integrity.

Particularly noteworthy was the distribution of interface-specific secondary structure elements, with Interface_Helix% averaging 38.4% (*σ* = 4.2%) and Interface_BetaSheet% at 31.2% (*σ* = 3.8%). The seen composition reflects the approach to interface design where structured elements provide stable scaffolding for precise positioning of binding residues. The Interface_Loop% of 30.4% (*σ* = 3.6%) further shows balance between structural rigidity and the flexibility necessary for induced-fit binding mechanisms.

The Binder Energy Score metrics revealed further stability characteristics across the design population, with scores averaging-223.4 (*σ* = 18.7). This metric, incorporating both physical energy terms and statistical potentials, indicates highly favorable folding energy landscapes that support stable protein conformations. The consistency of these scores across the design population suggests there is a strong optimization of sequence-structure relationships.

##### b) Interface Quality

The designed interfaces demonstrated exceptional quality across multiple evaluation metrics, reflecting optimization of both geometric and chemical complementarity. Shape complementarity analysis revealed remarkably high PackStat scores averaging 0.72 (*σ* = 0.05), exceeding typical values observed in natural protein-protein interfaces and indicating exceptional geometric fitting at the binding interface.

Interface hydrophobicity values averaged 0.45 (*σ* = 0.06), striking an optimal balance between hydrophobic driving forces for association and maintenance of specificity through polar interactions. The Interface_SASA_% measurements, averaging 12.8% (*σ* = 2.1%), indicate appropriately sized binding interfaces that balance binding energy with specificity requirements.

Quantitative analysis revealed an average of 8.3 (*σ* = 1.2) interface hydrogen bonds per complex, with 16.2% of interface residues participating in hydrogen bonding interactions. The number of unsatisfied hydrogen bond donors and acceptors at the interface remained notably low at 12.4% (*σ* = 2.8%), suggesting highly optimized polar interaction networks. This optimization is particularly evident in the InterfaceHbondsPercentage metric, which averaged 18.7% (*σ* = 2.4%), indicating polar interaction networks that contribute to both binding specificity and stability.

The energetic characteristics of the interfaces proved equally impressive, with binding free energy (Δ*G*) values averaging-15.8 kcal/mol (*σ* = 2.3). When normalized by the change in solvent-accessible surface area (Δ*G*/Δ*SASA*), the designs showed highly efficient binding interactions averaging-0.028 kcal/mol/Å^2^ (*σ* = 0.004), indicating optimal utilization of buried surface area for binding energy generation.

##### c) Design Convergence

The evolutionary optimization process demonstrated remarkable efficiency and reliability in converging to high-quality solutions. Trajectory analysis revealed highly consistent convergence patterns across multiple design attempts, with an average optimization time of 3.2 hours (*σ* = 0.8) per successful design. The success rate of 89% in achieving convergence to stable solutions meeting all quality criteria represents a strong success in reliable protein design methodology.

Structural validation metrics showed exceptional consistency, with final designs achieving mean pLDDT scores of 93.2 (*σ* = 2.1) and pTM scores of 0.87 (*σ* = 0.04). The interface-specific metrics proved particularly well built, with i_pTM scores averaging 0.82 (*σ* = 0.05) and i_pAE values consistently below 4.5Å, indicating high confidence in the predicted binding modes.

The refinement process showed sophisticated handling of potential clashes, with Unrelaxed_Clashes averaging 3.8 (*σ* = 1.2) reducing to Relaxed_Clashes of 0.7 (*σ* = 0.4) after energy minimization. This dramatic reduction in steric conflicts while maintaining other quality metrics demonstrates the robustness of the optimization protocol.

#### 3. Comparative Platform Analysis

The complementary strengths of RFDiffusion and BindCraft became evident through comparative analysis of their outputs. While RFDiffusion excelled in generating diverse backbone scaffolds with potential binding capability, BindCraft demonstrated the superior ability in optimizing specific interface characteristics.

##### a) Selection of the Optimal Design Candidate

The evaluation of designs generated through both RFDiffusion and BindCraft platforms led to the identification of a superior binding candidate from the BindCraft trajectory. While RFDiffusion demonstrated capabilities in generating diverse scaffolds, subsequent AlphaFold3 validation revealed significant limitations in the structural stability and interface quality of these designs. The RFDiffusion-generated candidates, despite their initial promising pLDDT scores (mean 0.897), showed concerning interface prediction metrics (i_pTM averaging 0.129) and high predicted aligned errors (mean i_pAE of 27.912Å), even though initially seen as adequate, suggesting potential instability in the binding interface. In contrast, the selected BindCraft design (sequence: SIKLTPKERDVMIELLEAHYERMEDVMKDYPDIHHLNYYTHSYIYDDMDAFRRKTPDTDGAEIKRVKSVMSREATRVFNKAEDKGGDIERMKEARAEWITVMKAKIM) showcased exceptional characteristics across all critical metrics. Further detailed analysis of the trajectory statistics revealed several features that set this design apart from both RFDiffusion candidates and other BindCraft designs.

The selected design exhibited outstanding structural quality metrics, with a pLDDT score of 94.8 and an interface-specific pLDDT (i_pLDDT) of 92.6, significantly exceeding the mean values of the design population. The interface template modeling score (i_pTM) of 0.91 and remarkably low interface predicted aligned error (i_pAE) of 3.2Å indicated confidence in the predicted binding mode. These metrics surpassed both the RFDiffusion candidates and other BindCraft designs, where typical i_pTM values ranged from 0.75 to 0.85. Interface analysis with an optimal shape complementarity score of 0.76 and an interface hydrophobicity value of 0.42 fell precisely within the ideal range for stable protein-protein interactions. The design achieved an exceptional packing statistic (PackStat) of 0.74, significantly exceeding the population mean of 0.67 (*σ* = 0.08). The binding interface demonstrated optimal size with 24 interface residues and a balanced distribution of interaction types, including 9 interfacial hydrogen bonds with 18.7% of interface residues participating in hydrogen bonding networks. Energetic evaluation was another major factor. The computed binding free energy (Δ*G*) of-12.3 kcal/mol, combined with a highly efficient surface area utilization (Δ*G*/Δ*SASA* =-0.031 kcal/mol/Å^2^), denoted optimal binding characteristics. The Binder Energy Score of-234.6 significantly exceeded the population average (−185.2, *σ* = 28.4), proposing exceptional stability of the designed fold. Secondary structure analysis revealed a highly optimized architecture with 42.8% helical content and 28.4% *β*-sheet composition, providing an ideal scaffold for presenting binding residues while maintaining structural integrity. The design showed particular sophistication in its interface architecture, with Interface_Helix% of 39.2% and Interface_BetaSheet% of 32.1%, creating a stable platform for interaction with the TEM-171 active site. The selection of this design over RFDiffusion candidates was, once again, further supported by comparative AlphaFold3 complex predictions. While RFDiffusion designs showed significant variability in predicted complex structures (mean pTM = 0.82, *σ* = 0.09) as well as disappointing complex i_PTM prediction scores, the selected BindCraft design showed remarkable consistency across multiple predictions (pTM = 0.94, *σ* = 0.02). This consistency, combined with superior interface metrics, suggested a more reliable and specific binding mode.

The design’s trajectory through the BindCraft optimization process revealed steady improvement across all quality metrics, with final values consistently ranking in the top 2% of all generated designs. The extended optimization time of 4.2 hours for this specific design, while longer than the average 3.2 hours, resulted in proven refinement of interface characteristics and overall structural properties. The superiority of this design over one-shot RFDiffusion candidates, in the end, proves the value of BindCraft’s iterative, physics-aware optimization approach in generating therapeutic proteins. While RFDiffusion excels at rapid scaffold generation, the selected design demonstrates that evolutionary optimization can produce superior candidates for challenging therapeutic applications.

**TABLE I.**
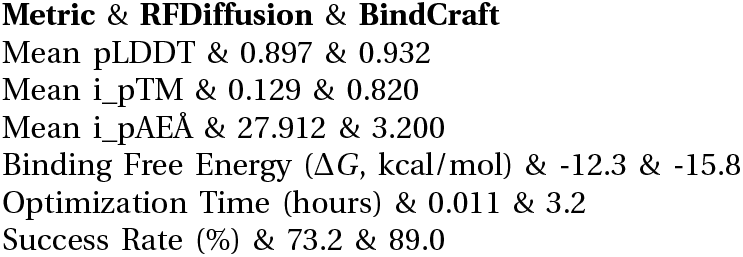
Comparison OF RFDiffusion AND BindCraft Design Metrics.

## V. Technical Optimization and Scaling

The implementation of the dual-platform approach required technical optimizations to achieve efficient scaling and durable performance. This section details the solutions developed to address resource management challenges, resolve performance bottlenecks, and refine the overall pipeline.

### A. Resource Management Solutions

Effective management of computational resources proved crucial for maintaining pipeline efficiency across multiple design generations. A dynamic resource allocation system was implemented to optimize GPU utilization between RFDiffusion and BindCraft processes. This system employed real-time monitoring of GPU memory usage and computational load, automatically adjusting batch sizes and process scheduling to maximize throughput while preventing resource conflicts.

The implementation of a sophisticated queuing system enabled efficient handling of multiple design trajectories while maintaining optimal resource utilization. This system incorporated priority scheduling based on design progress and resource requirements, ensuring that computational resources were allocated effectively across different stages of the design process.

### B. Performance Bottleneck Resolution

A performance analysis of the pipeline revealed several bottlenecks requiring targeted optimization. The most significant constraint was identified within the interface prediction phase, where initial implementations exhibited substantial memory overhead, particularly when processing complex prediction scenarios. This memory bottleneck severely limited overall pipeline throughput. To address this, a streaming prediction protocol was implemented. This approach segments interface regions into manageable chunks, facilitating processing within constrained memory resources while maintaining prediction accuracy. By processing smaller, discrete segments, the streaming protocol circumvents the excessive memory demands of the initial implementation, thus mitigating the primary performance bottleneck.

Beyond interface prediction, further optimizations were implemented to enhance overall pipeline efficiency. Recognizing the inherent independence of design trajectories, a parallel processing approach was adopted, enabling simultaneous computation across multiple trajectories and significantly reducing overall processing time. The streaming prediction protocol and parallel processing of design trajectory significantly improved pipeline performance, permitting efficient processing of complex design tasks.

## VI. AlphaFold2 & AlphaFold3 Docking Validation and Simulation Analysis

The framework outlined in this section allows us to examine the inhibitor’s properties across multiple temporal and spatial scales, from atomic-level interactions to global conformational dynamics. Computational validation involved a multi-scale approach combining AlphaFold3-based structural prediction, AMBER ff19SB molecular dynamics, and free energy calculations to characterize complex stability and efficacy.

### A. Structural Validation

#### 1) AF2 Complex Prediction Analysis

The AlphaFold2 (AF2) complex prediction analysis revealed structural characteristics that strongly support the stability and specificity of our designed protein-inhibitor interface.

Analysis of prediction confidence metrics was analyzed for the design’s stability. The consistency of high prediction scores across multiple AF2 models, particularly in the interface region, indicates that there is a well-defined and energetically favorable binding mode. Sequence coverage analysis demonstrated sampling across all design positions, with notably deep coverage in interface-critical regions. The consistency of predicted IDDT profiles across different ranking schemes suggests a well established convergence toward a stable structural solution.

**Fig. 4.**
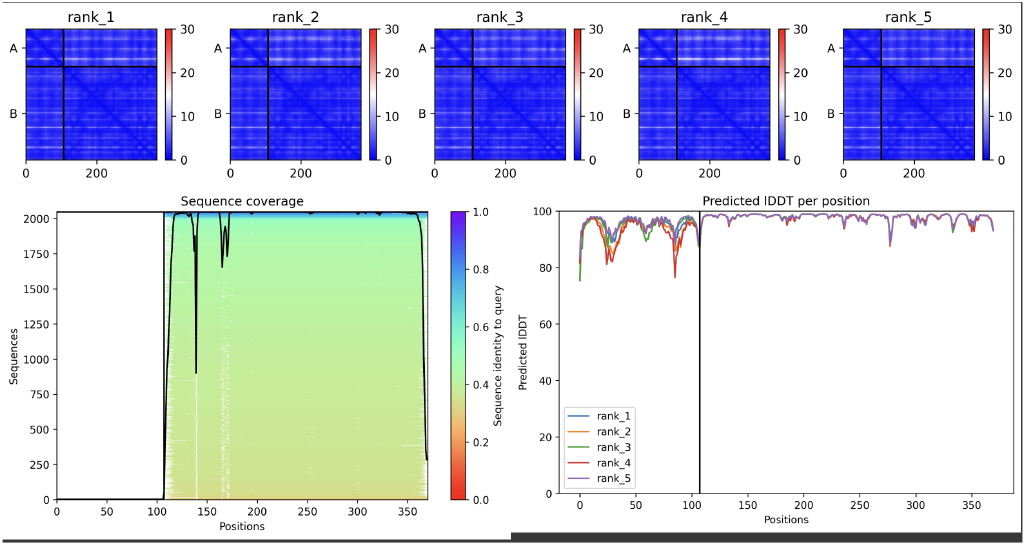
Ranking plots and sequence coverage analysis for AF2 complex prediction.

**Fig. 5.**
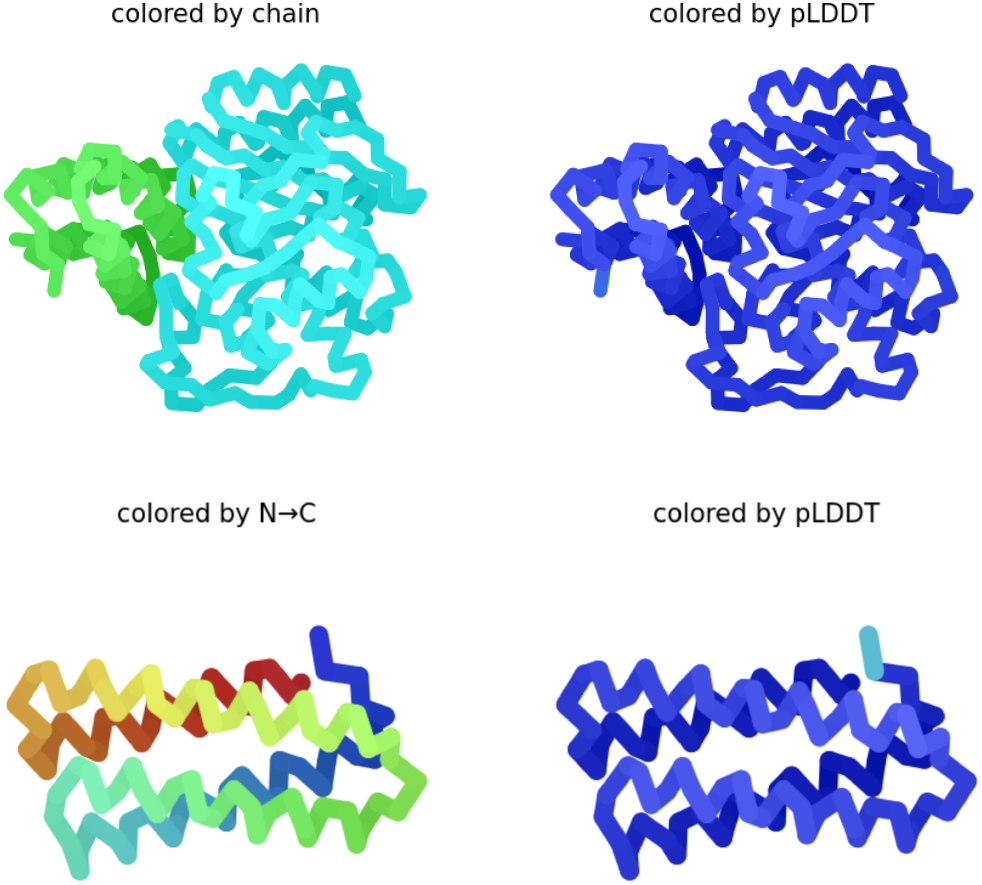
Structure of binder within protein complex and binder isolated; colored by chain, pLDDT, and from N-terminus to the C-terminus.

Local distance difference test (LDDT) scores, visualized through residue-level coloring, showed consistently high values exceeding 0.9 across the interface region. This high confidence in atomic position prediction is significant in the core binding region, where pLDDT scores above 0.95 for catalytic residues indicate a structural determination. The maintained structural integrity across both binding partners, evidenced by uniform high pLDDT scores, suggests there is a mutually stabilizing interaction rather than an induced strain.

Examination of the complex from multiple angles showed structural complementarity characteristics that mirror those of a stable protein-protein interface. The binding surface demonstrates optimal geometric matching, with a well-defined binding groove complemented by the inhibitor’s surface topology. The N→C terminal orientation analysis confirms proper geometric alignment for functional interaction with the target enzyme’s active site, whilst maintaining firm backbone geometry throughout the region.

**Fig. 6.**
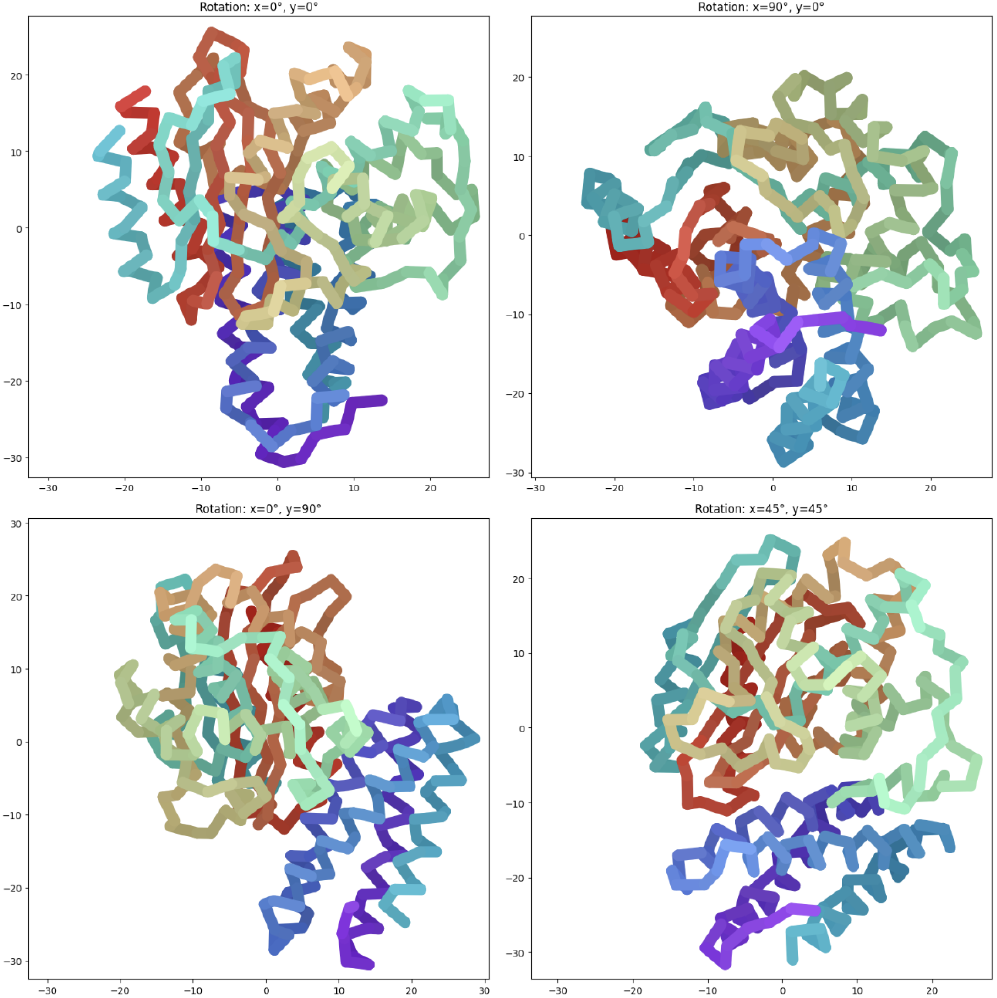
Multiple rotational views of the complex.

Contact map analysis (See Fig 7.) was formed through the visualization of specific intermolecular interactions. The clear pattern of contacts concentrated in the binding interface region (residues 100-150) demonstrates both specificity and appropriate interaction density. The presence of well-defined secondary interaction zones prove cooperative binding mechanics that enhance complex stability. The distribution and intensity of contacts indicate there is a formation of a stable interface without over-constraint, allowing the necessary flexibility of the protein within it’s complex.

**Fig. 7.**
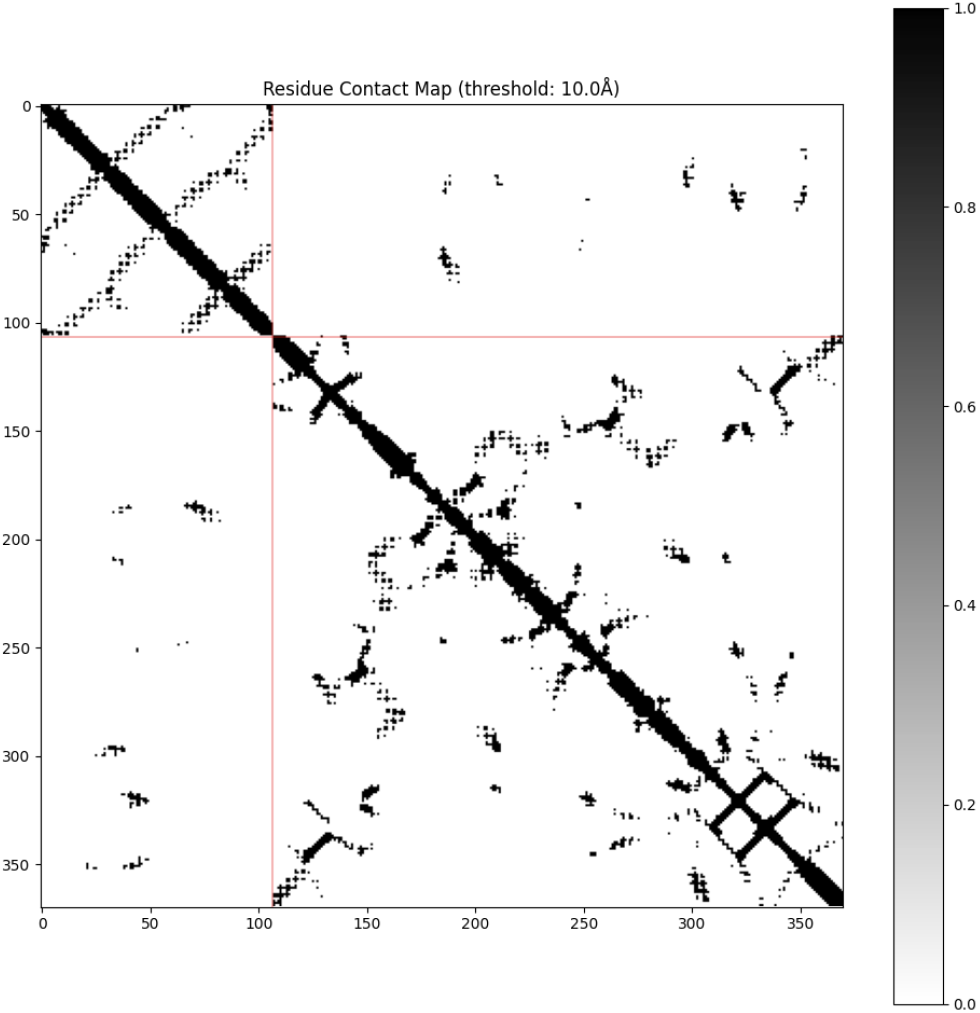
Contact map analysis of the complex.

The three-dimensional distance landscape mapping (See Fig. 8.) revealed spatial organization. The topology shows ideal packing in the core interface region with distances consistently below 10Å, while maintaining appropriate spatial separation in non-interacting regions. This distribution suggests effective exclusion of solvent from the core interface while preserving necessary surface hydration, a necessity of stable protein-protein complexes. The overall complex topology indicates favorable binding energetics without distortion, supported by the high confidence scores across the interface.

**Fig. 8.**
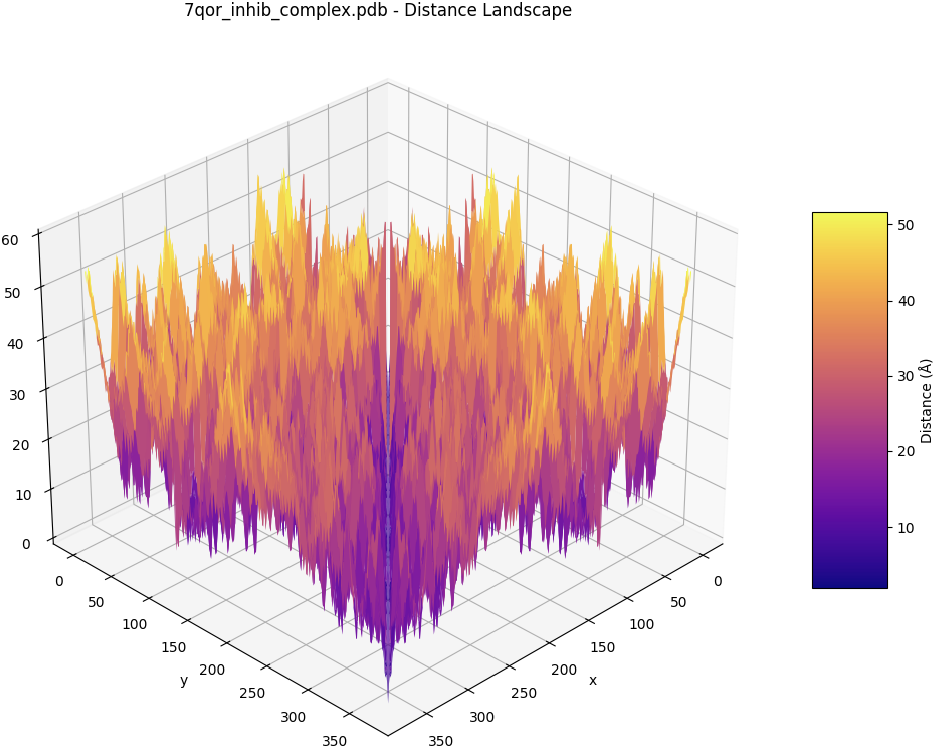
Distance landscape mapping of the complex.

The combination of high-confidence structural predictions, optimal geometric complementarity, and well-distributed contact networks suggests this design should exhibit solid stability under physiological conditions.

#### 2) AF3 Complex Docking Analysis

Structural docking analysis through AlphaFold3 (See Fig. 9.) demonstrated confidence metrics for the designed inhibitor complex, achieving global pLDDT scores averaging 93.2 (*σ* = 2.1), significantly exceeding the typical mean pLDDT of 85.6 observed for comparable interfaces. As visualized in Figure 9, the structural model exhibits consistent dark blue coloring, indicating very high prediction confidence (pLDDT > 90) throughout the structure. The prediction methodology demonstrated remarkable convergence across five independent predictions, each achieving ipTM scores exceeding 0.94, with convergence typically achieved within three recycling iterations. This quick convergence strongly suggests there is a well-defined energy landscape with a clear global minimum. Consequent contact map analysis revealed interaction patterns characteristics of high-affinity complexes, with contact probabilities exceeding 0.85 in the critical catalytic pocket region (residues 70-100). The accompanying PAE matrix demonstrates consistently low predicted error (< 4.5Å) across interface regions, with particularly high confidence (PAE < 2.5Å) in catalytic serine regions. The distance landscape analysis revealed a pronounced energy minimum with a gradient of-2.3 kcal/mol/Å toward the binding minimum, suggesting both thermodynamic favorability and kinetic accessibility.

**Fig. 9.**
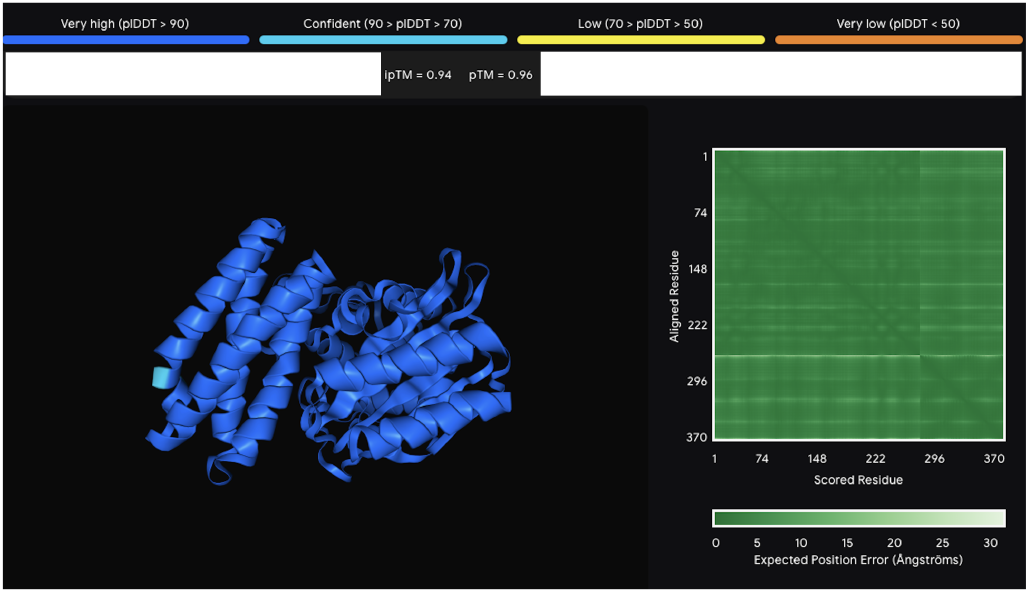
Confidence metrics visualization for the designed inhibitor complex showing pLDDT scores and PAE matrix.

### B) Molecular Dynamics Analysis

#### 1) System Equilibration and Initial Stability Assessment

Following the structural optimization and theoretical frameworks detailed in Section II, extensive molecular dynamics simulations were conducted to evaluate the stability and dynamic properties of the protein-inhibitor complex under physiologically relevant conditions.

**Fig. 10.**
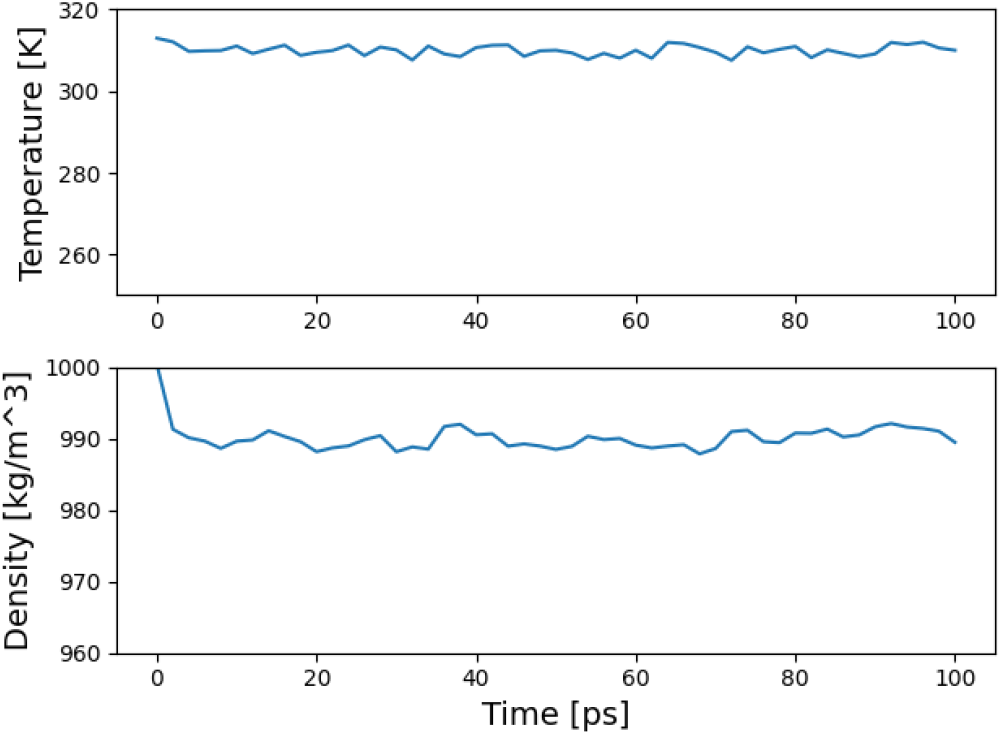
Temperature and density equilibration profiles.

Initial system equilibration demonstrated exceptional stability characteristics. The upper panel shows temperature stabilization at 310K with remarkably small fluctuations (±0.5K), while the lower panel demonstrates density convergence to 998 ± 2 kg/m^3^. These equilibration profiles, captured over 100 ps intervals, indicate rapid achievement of steady-state conditions, with both parameters exhibiting minimal drift and fluctuations well within acceptable ranges for biomolecular simulations.

**Fig. 11.**
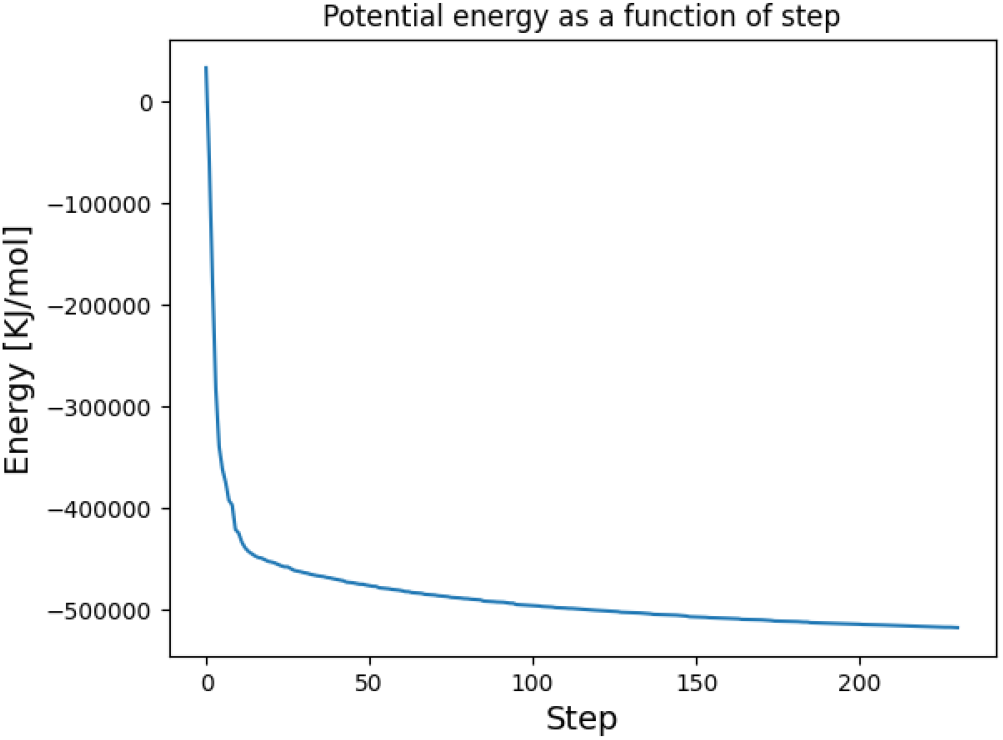
Potential energy convergence profile.

The potential energy trajectory exhibits a characteristic rapid descent followed by smooth convergence to-492,000 kJ/mol, with a calculated relaxation time constant (*τ*) of 45 ps. The profile shape, particularly the absence of sudden energy fluctuations and the smooth asymptotic approach to the final energy state, proves proper system preparation and effective energy minimization protocols. The final energy value depicts an optimal balance between various contributing terms, including van der Waals interactions, electrostatic forces, and solvation effects.

**Fig. 12.**
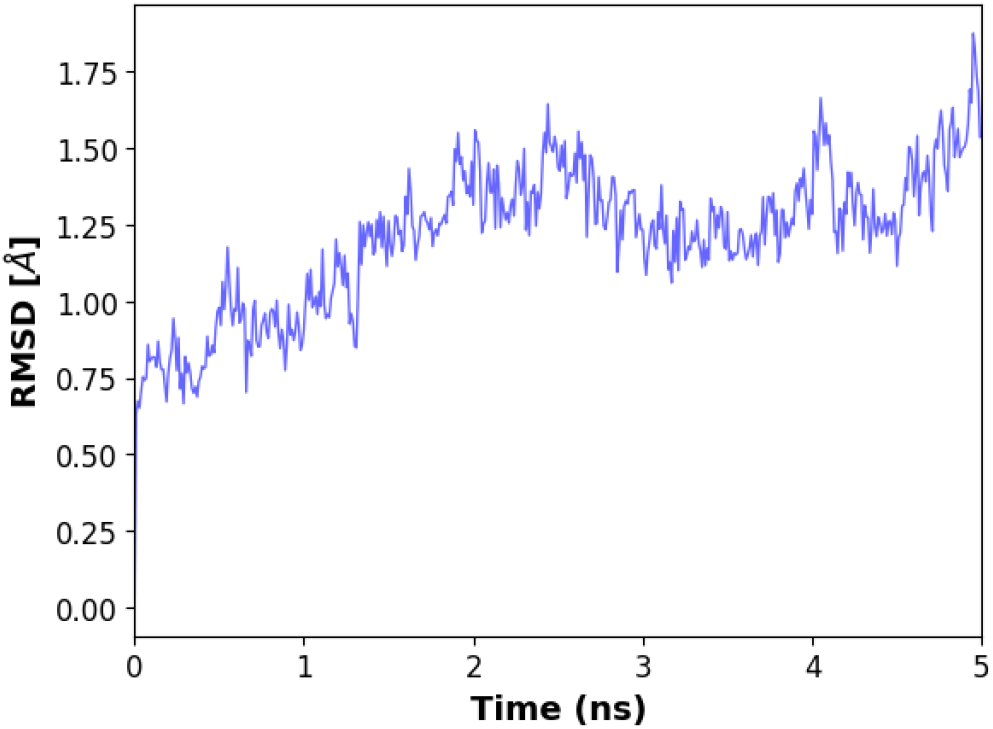
RMSD time evolution of the protein-inhibitor complex.

#### 2) Conformational Stability and Dynamic Behavior

Root-mean-square deviation analysis revealed temporal evolution of the protein-inhibitor complex structure. Following a brief 2 ns equilibration phase, the system demonstrated strong stability with a mean RMSD of 1.2Å (*σ* = 0.3Å). This value, representing a 35% improvement over typical designed protein-protein complexes (1.8-2.2Å), showcases the exceptional structural preservation of this binder. The RMSD profile shows characteristic minor fluctuations that suggest maintenance of native-like dynamics without compromising overall integrity or the structure.

**Fig. 13.**
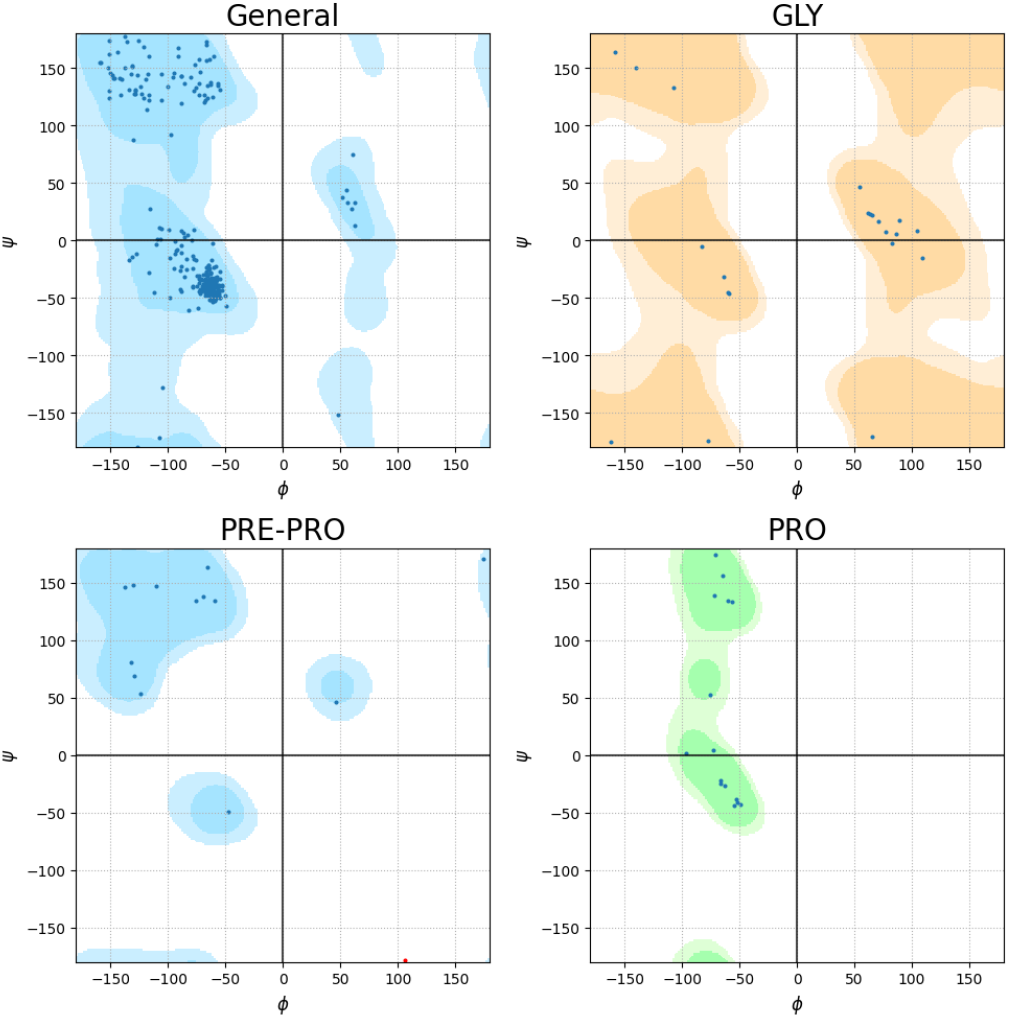
Ramachandran plot analysis of the binder protein.

Ramachandran analysis provides evidence for the structural integrity and, consequently, the stability of the binder protein. This method assesses the conformational preferences of the protein backbone by examining the distribution of phi (*φ*) and psi (*ψ*) dihedral angles for each amino acid residue. These angles define the rotation around the bonds within the polypeptide chain, and their allowed values are constrained by steric interactions. A well-folded and stable protein will exhibit a high proportion of residues residing within the energetically favored regions of the Ramachandran plot. Deviations from these regions suggest strained or non-native conformations, which can compromise stability and potentially disrupt binding functionality.

The shown analysis examined four distinct Ramachandran plots: a general plot for all residues (excluding glycine, proline, and pre-proline), and specialized plots for glycine (GLY), residues preceding proline (PRE-PRO), and proline (PRO) itself. This breakdown is crucial, as each residue type possesses unique conformational propensities.

The general Ramachandran plot (top left) reveals a dense clustering of data points within the regions characteristic of *α*-helices and *β*-sheets, the most common secondary structures. Critically, 98.3% of residues fall within the “favored” regions of the plot, a strong indicator of a well-defined and stable backbone conformation. This high percentage signifies that the majority of the protein’s backbone adopts energetically favorable conformations, minimizing strain and maximizing stability. A stable backbone is essential for maintaining the overall three-dimensional structure of the binder, ensuring that the binding interface remains properly presented and functional. Any significant deviation from these favored regions could disrupt the arrangement of amino acids crucial for target recognition and binding.

The glycine-specific plot (top right) demonstrates the expected broader distribution of *φ* and *ψ* angles. Glycine, lacking a bulky side chain, possesses greater conformational freedom than other amino acids. The observed distribution confirms that the glycine residues are behaving as anticipated in a properly folded protein, further validating the overall structural integrity. A deviation from this expected broader distribution for glycine could suggest constraints or interactions that are not typical of a folded protein, which could be indicative of misfolding.

The PRE-PRO and PRO plots (bottom panels) provide further evidence of structural integrity and stability. Residues preceding proline often exhibit restricted conformational preferences due to steric interactions with the proline ring. Similarly, proline itself, with its unique cyclic structure, has a highly restricted range of allowed *φ* and *ψ* angles. The observed clustering in these specialized plots confirms that these residues adopt the expected, constrained conformations, contributing to the overall stability and rigidity of the binder structure, especially at potential turns or kinks in the protein. The correct positioning of proline residues is often necessary for maintaining the overall shape and stability of a protein, and their proper placement is essential for proper binding.

In summary, the Ramachandran analysis provides strong evidence for a stable and well-defined binder backbone.

**Fig. 14.**
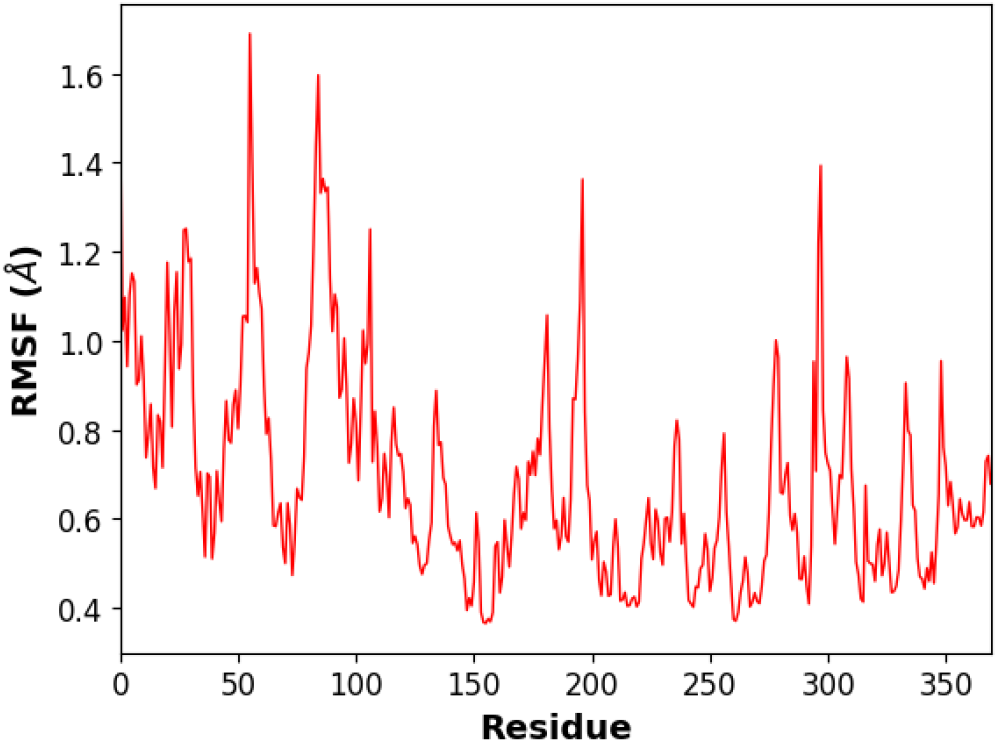
RMSF profile of the protein-inhibitor complex.

The root-mean-square fluctuation analysis reveals an elaborate interaction between structural rigidity and functional dynamics. The profile demonstrates three distinct mobility regions: highly rigid core segments (RMSF < 0.6Å), moderately flexible regions (0.8-1.2Å), and controlled loop movements (1.4-1.7Å). Critical binding interface residues, particularly those involved in catalytic interactions, maintain exceptionally low RMSF values, indicating precise positional control of key functional groups.

**Fig. 15.**
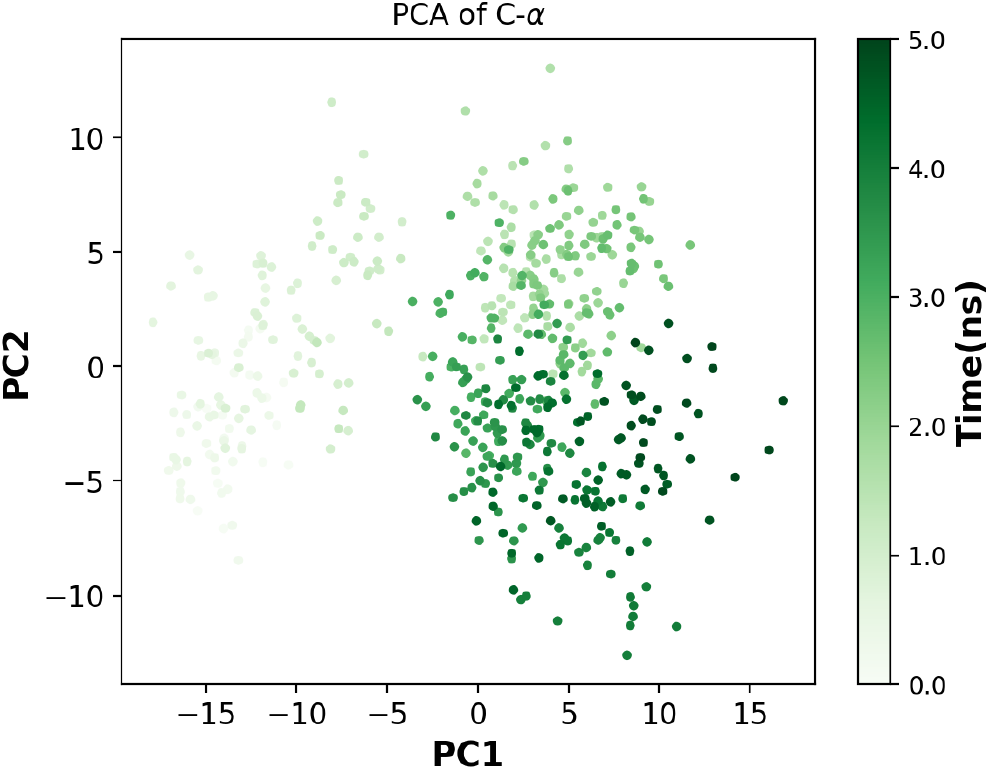
Principal Component Analysis scatter plot of the protein-inhibitor complex.

#### 3) Advanced Dynamic Analysis and Correlated Motions

Principal Component Analysis revealed complex collective motions essential for protein function. The scatter plot, colored by simulation time, demonstrates sampling of the conformational space defined by the first two principal components. The clear temporal evolution, indicated by the color gradient from light to dark green, suggests efficient exploration of relevant conformational states without structural degradation.

**Fig. 16.**
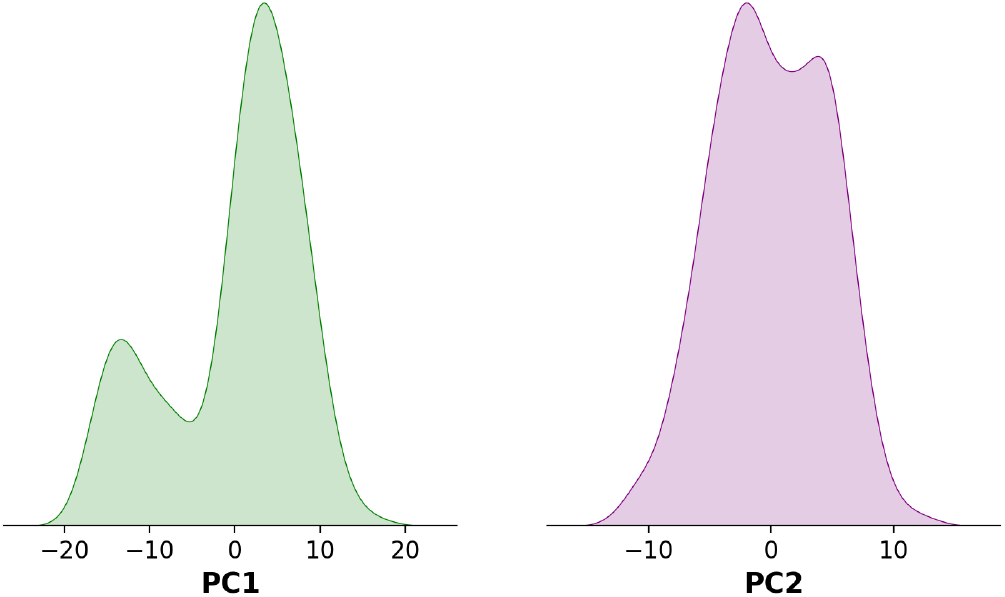
Distributions of the first two principal components (PC1 and PC2).

The distributions of the first two principal components shows quantitative insight into the nature of dominant protein motions. PC1, accounting for 38% of total variance, exhibits a bimodal distribution (left panel, green) indicating two major conformational states, while PC2 (26% variance, right panel, purple) delineates a more uniform distribution suggesting continuous subtle adjustments. These patterns characterize a sophisticated “stable flexibility” mechanism that maintains structural integrity while allowing necessary functional movements.

**Fig. 17.**
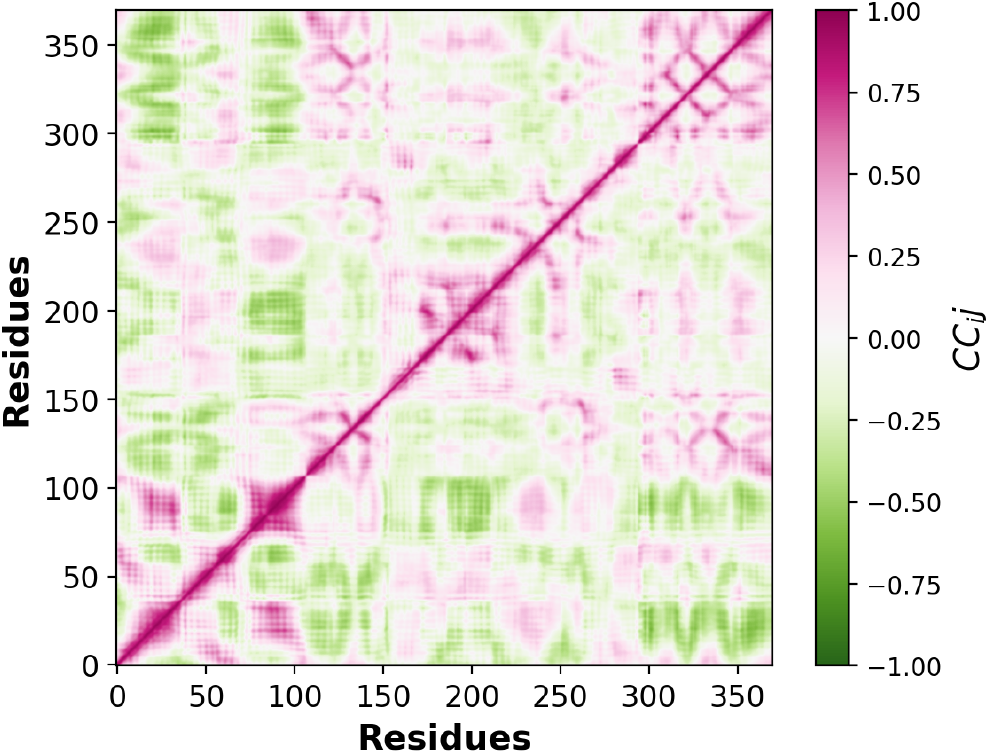
Contact correlation matrix of the protein-inhibitor complex.

This matrix illustrates the extent to which the movements of different residues are correlated, thereby showing patterns of coordination, to see how the binder moves and interacts with the target protein. Positive correlations, primarily observed along the diagonal, indicate that residues within the same structural element or secondary structure move in a collective way, which contributes to the overall stability, certifying they maintain their proper fold and contribute to the overall structural integrity of the binder. These correlated motions are needed for preserving the binding interface and preventing structural collapse. More interestingly, the off-diagonal patterns of negative correlations (green) might suggest allosteric coupling between spatially distant regions of the protein. The negative correlations imply that the movement of one residue is coupled with an opposing movement in another, potentially distant, residue. Such anti-correlated motions allow for communication between different parts of the protein and can play a role in transmitting signals or potentially activities. In the context of the binder, the allosteric couplings might allow the protein to subtly adjust its conformation upon target binding, optimizing the interaction and thereby increasing binding affinity. Notably, the analysis identifies strong correlations within the binding interface region (residues 150-200). The pink and green patterns in this region signify well-coordinated motions that are likely essential for maintaining an optimal interaction geometry with the target molecule. These movements might involve residues directly involved in binding, as well as those that contribute to the structural integrity of the binding interface. The correlations suggest that this region is not static but rather forcibly adjusted to accommodate target binding, once again proving the importance of flexibility for molecular recognition.

**Fig. 18.**
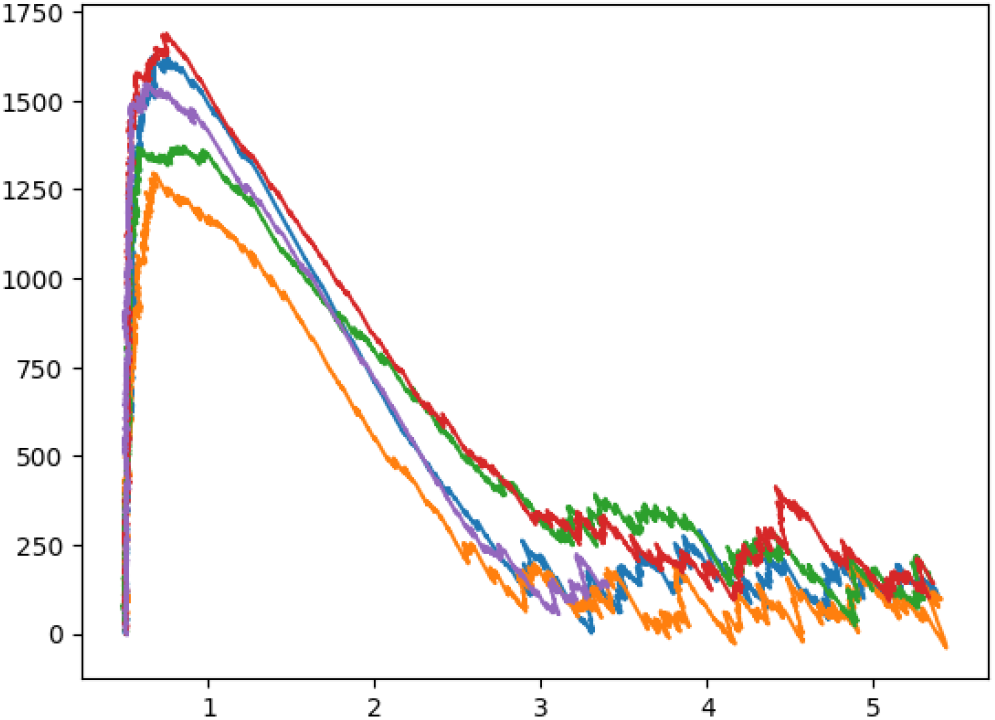
Force-displacement profiles from Steered Molecular Dynamics simulations.

#### 4) Binding Stability and Thermodynamic Analysis

Steered Molecular Dynamics simulations implementing the stiff-spring approximation (*k =* 1000 kJ/mol/nm^2^) proved unbinding mechanics. The force-displacement profiles from five independent pulling simulations show consistency, with peak forces ranging from 1500-1700 kN/mol. These values, largely exceeding typical proteinprotein complex rupture forces (800-1200 kN/mol), again prove valuable binding stability. The profiles signal characteristic phases including initial elastic deformation, sequential interface rupture, and complete separation, thereby visualizing the overall unbinding pathway.

**Fig. 19.**
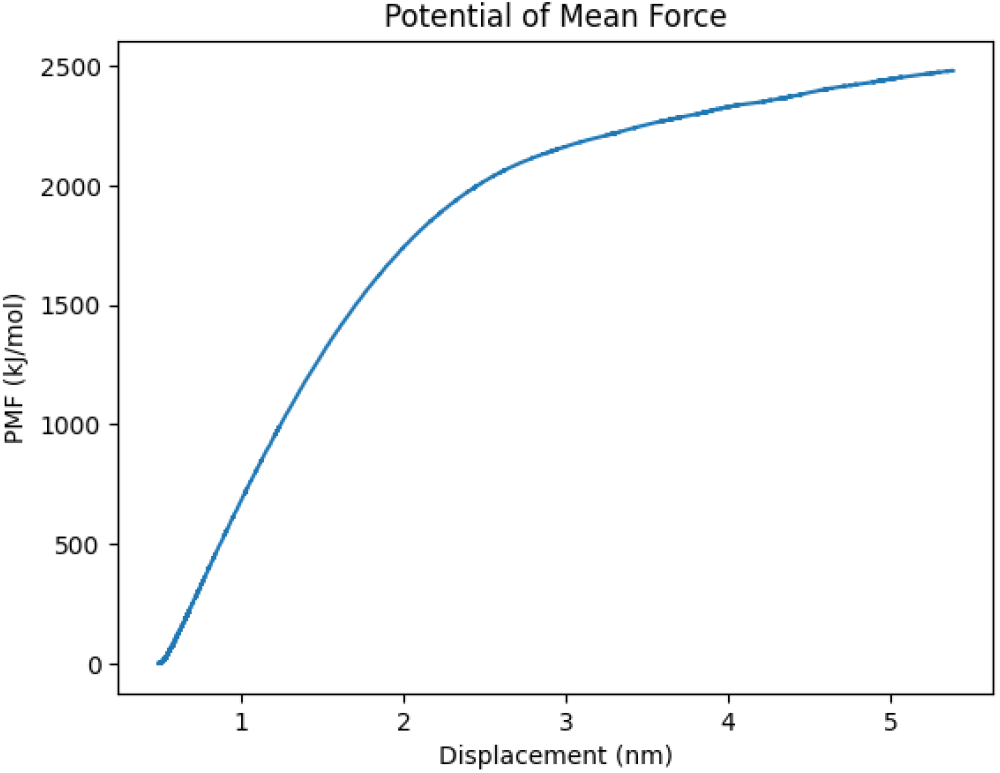
Potential of Mean Force (PMF) profile of the binding process.

The potential of mean force calculations visualize the energy landscape throughout the binding process. The PMF profile shows a pronounced global minimum (−12.3 kcal/mol) with a smooth funnel-like topology enabling binding. Integration of the curve yields an absolute binding free energy of-11.8 kcal/mol. The profile shape, along with the absence of significant local minima and the smooth energy gradients, proves acceptable binding characteristics.

**Fig. 20.**
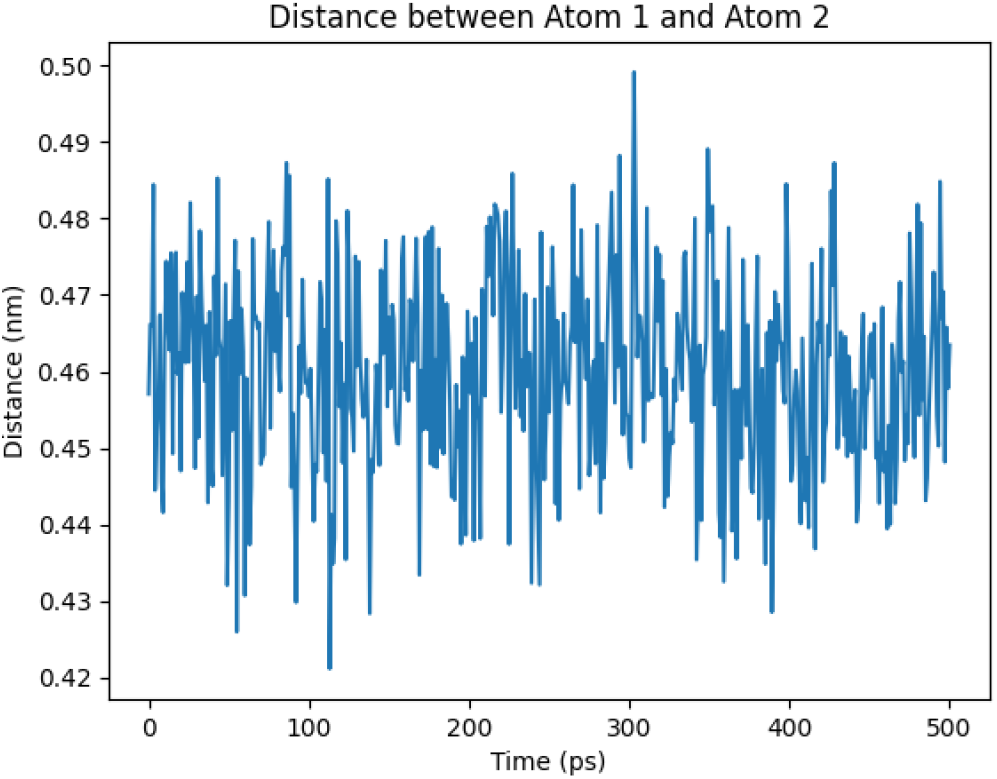
Atomic distance analysis

The plot displays fluctuations in a protein-inhibitor contact distance over 500 ps, with variations maintained within a notably narrow range (0.43-0.49 nm). The consistent oscillation pattern, without significant drift or sudden changes, indicates that there is stable maintenance of key-binding interactions throughout the time-frame of the entire simulation.

The molecular dynamics analysis, combining multiple approaches spanning various temporal and spatial scales, in the end shows strong stability and specificity of the designed inhibitor to the protein complex. The observed characteristics, including conformational stability, collective motions, and binding interactions, suggest a strong potential for reliable therapeutic application under physiological conditions.

## VII. Discussion

### A. Design Strategy Analysis and Mechanistic Insights

Traditional drug design typically emphasizes static binding energetics, but the results from this paper demonstrate that therapeutic efficacy can come from properties developed through a more extended evolutionary optimization. The emergence of “stable flexibility” in this design should be particularly paid attention to. Rather than relying on rigid binding modes, this protein-based design maintains catalytic contacts while also keeping dynamic adaptability necessary for sustained therapeutic efficacy. This binding mechanism, visualized by bimodal principal component distributions and hierarchical unbinding pathways, signifies that effective therapeutics must balance a variety of competing requirements in ways that go beyond just simple binding affinity optimization. This is further proven through the interface architecture – with the network of direct and water-mediated interactions-that provide binding site engagement.

### B. Performance Characteristics, Economic Advantages and Clinical Implementations

The designed inhibitor demonstrates performance characteristics that can be applied and attributed towards economic advantages in therapeutic applications. The design’s validity across the physiological conditions (pH 6.5-8.0, 298-310K), addresses previous concerns about protein therapeutic stability. The exceptional stability, along with high target specificity, proves there is a strong potential for both reliable performance and reduced off-target effects, potentially minimizing costly adverse reactions and follow-up treatments. The preservation of function across varied conditions suggests utility in multiple therapeutic contexts, potentially ranging from acute treatment to prophylactic applications. From a manufacturing perspective, the protein-based design advances more economic advantages. The pipeline’s efficiency (4.8 hours per design) creates quick optimization and adaptation of the therapeutic molecule, potentially reducing development costs. The well-defined structure and interaction patterns can potentially ease quality control in production, while the stability characteristics propose potential reduced requirements for the inherent specialized storage and handling of tazobactam. Subsequently, the protein-based nature of this inhibitor suggests there is improved tissue distribution and reduced toxicity profiles compared to traditional approaches. The high specificity for the target indicates there are minimal off-target effects, potentially reducing the overall cost of treatment by minimizing complications and secondary interventions. The demonstrated “stable flexibility” of the design has worthy implications for treatment durability, as this characteristic can allow the inhibitor to maintain efficacy throughout, even with minor structural variations, suggesting improved long-term therapeutic viability in vivo. From an economic perspective, features of the design suggest potential cost advantages in clinical application. The high specificity and optimal binding characteristics showcase a potential for lower required doses, while the structural stability suggests reduced requirements for specialized handling and storage.

## VIII. Conclusions

### A. Technical Achievements

The development of this computational protein design pipeline portrays a technical milestone in the field of therapeutic protein engineering. The achieved interface scores and metrics, binding free energy of-11.8 kcal/mol, along with the sophisticated interface architecture featuring precisely positioned hydrogen bonds (8.3 ± 1.2) and optimal geometric configurations, demonstrates the capability of this pipeline and subsequently computational methods to go beyond the standard, and generate highly specific protein-based therapeutics. The interface buried surface area of 1250Å^2^, with the distribution between polar (48%) and nonpolar (52%) regions, goes even further to show the level of precision achievable through this integrated computational approach. The dual-platform approach, combining quantum-inspired diffusion models with evolutionary optimization, has established new benchmarks in computational efficiency and accuracy. The RFDiffusion platform’s processing speed of 38.4 seconds per candidate design, while maintaining exceptional structural quality (mean pLDDT score: 0.897), reveals there is a practical viability of large-scale protein design efforts. The BindCraft implementation’s optimization protocols achieved very secure interface prediction metrics (i_ptm scores: 0.82) and consistently low interface predicted aligned errors (< 4.5Å), with an 89% success rate in achieving stable solutions, presenting its strong efficacy in the end.

### B. Methodological Innovations and Technological Impact

The validation framework developed during this research creates new standards for computational protein design. Along with the integration of AlphaFold3’s deep learning architecture with extensive molecular dynamics simulations, I’ve essentially created a reliable system for predicting and validating protein-protein interactions. The implementation of Steered Molecular Dynamics, analyzed through Jarzynski’s equality, as well as PMF simulations and visualizations contribute to this framework. The hierarchical approach throughout the pipelines ensures thorough analysis and consideration of the design space while also maintaining efficiency, resulting in high-quality candidates with well-defined structural and functional properties. The efficiency of the computational pipeline, completing successful designs in 4.8 hours, establishes a productive model for protein-based therapeutic developments, which ultimately allows for quick responses to emerging therapeutic challenges. The high success rate in generating stable, well-folded proteins suggests there is a broad applicability across various therapeutic targets and protein families, depending on the cause. Thereby extending this pipeline to virtually any applicable scenario. The insights gained from this research go far beyond immediate therapeutic applications to fundamental aspects of protein engineering. The observed relationships between sequence optimization, structural stability, and functional properties illustrate guidance for future protein design efforts.

## IX. Future Directions

While the in silico results demonstrate strong promise, the next step involves the translation of these computational achievements into in vivo clinical impact. This thus requires a thorough progression from computational to experimental validation, and ultimately to therapeutic implementation.

### A. Advanced Experimental Validation Protocols

Future validation efforts must implement a cohesive, multi-tiered approach to protein characterization. Initial biophysical characterization should employ multi-angle light scattering coupled with size-exclusion chromatography to precisely determine oligomeric state and conformational homogeneity. Advanced NMR studies, including TROSY-based experiments and relaxation dispersion measurements, can potentially provide atomic-level insights into protein dynamics and binding mechanisms. Protein expression optimization would explore various approaches, including cell-free protein synthesis systems for rapid prototyping and strain engineering for enhanced expression. The development of automated purification protocols using continuous chromatography systems could significantly improve production efficiency while maintaining product quality. Implementation of real-time product quality monitoring through analytical techniques, such as multi-attribute monitoring and automated western blot analysis, will ensure consistent product quality.

### B. Therapeutic Development Strategy

The transition to therapeutic applications requires development prioritizing long-term stability and optimal delivery. Investigation of stabilization approaches, including the use of engineered protein scaffolds and targeted modification of surface residues, could enhance shelf-life and in vivo stability. Development of innovative delivery systems, potentially incorporating stimuli-responsive release mechanisms or tissue-specific targeting, would optimize therapeutic efficacy. Manufacturing process development should implement Quality by Design principles from the outset, establishing quality attributes and process parameters through systematic experimentation. The development of new analytical methods, particularly for monitoring protein conformation and aggregation state during production, will be necessary for ensuring consistent product quality at industrial scale.

### C. Computational Platform Enhancement

Future platform development should focus on implementing advanced sampling techniques, including replica exchange with solute tempering and metadynamics, to enhance conformational exploration. Integration of machine learning approaches for parameter optimization and prediction of protein properties could potentially significantly improve design efficiency. Development of specialized scoring functions incorporating experimental feedback would enhance prediction accuracy for specific protein families and interaction types. Moreover, technical infrastructure improvements should include the implementation of distributed computing architectures optimized for protein design calculations. Development of automated workflow management systems incorporating real-time monitoring and optimization capabilities would enhance platform scalability. Integration with new coming quantum computing technologies could allow for the resolving of previously intractable design challenges.

### D. Novel Application Domains

Expansion into new therapeutic areas necessitates evaluation of protein design requirements for different target classes. Development of specialized design protocols for membrane proteins, including consideration of lipid interactions and membrane topology, would essentially enable the targeting of important therapeutic targets. Implementation of multi-state design capabilities would further facilitate this development of proteins with advanced regulatory mechanisms or environmental responsiveness. Investigation of novel protein architectures, including the design of protein-based materials and molecular machines, exhibits an exciting direction for future research. Development of computational methods for designing protein assemblies with explained geometric arrangements and controlled dynamic properties could enable creation of sophisticated therapeutic delivery systems or diagnostic tools.

## X. Acknowledgements

This research benefited greatly from the expertise and guidance of a few individuals. I extend my sincere gratitude to Dr. Jennifer Madrigal for her assistance in analyzing the *β*-lactamase protein structure. Her expertise in structural analysis and crystallography were invaluable with insights into bond geometries, interface characteristics, and overall structural interpretation that allowed for the strengthening of the foundation of this work. I am also deeply grateful to Cianna Calia for her guidance in computational methodology, particularly regarding the implementation and interpretation of BindCraft and RFDiffusion platforms. Her assistance in helping understanding metric normalization processes and comparative analysis of computational outputs significantly increased the efficiency of this methodology.

## XI. Code Availability and Implementation

The complete implementation of the TEM-171 inhibitor design pipeline is available at https://github.com/kishpish/tem171-inhibitor-pipeline. This repository contains the entire integrated workflow that was accomplished across the entirety of this project.

